# Biomimetic miR-133a inhibitor activated scaffolds optimised for spinal cord repair promote neurite outgrowth and angiogenesis via neuronal cytoskeletal remodelling

**DOI:** 10.64898/2026.04.22.719922

**Authors:** Cian O’Connor, Rena Mullally, Juan Carlos Palomeque-Chávez, Marko Dobricic, Eoin McCoy, Jack Maughan, Rachel Stewart, Chayanika Saha, Julia O’Sullivan, Helen O. McCarthy, Maeve A. Caldwell, Jochen H.M. Prehn, Fergal J. O’Brien

## Abstract

Significant challenges in spinal cord injury include the loss of neural tissue, disruption of local vasculature, and intrinsic suppression of actin mobilisation in neurons, together preventing axonal regrowth. Here, we develop an implantable biomimetic microRNA (miR) inhibitor-activated scaffold that combines optimised matrix cues with transcriptomically defined RNA-based modulation of intrinsic neuronal pathways as a platform to support neuronal cell delivery and promote neurovascular repair. First, hyaluronic acid macroporous scaffolds functionalized with collagen-IV and fibronectin supported iPSC-derived neuronal spheroid formation and neurite extension. To identify a neurotrophic target, we performed analysis of public miRNA-mRNA interaction datasets, revealing that miR-133a regulates pathways involved in neuronal actin cytoskeletal organisation. MiR-133a inhibitors were complexed with the cell-penetrating peptide RALA to form nanoparticles, demonstrated >95% scaffold loading efficiency, sustained localised release over 28 days and enhanced neurite outgrowth from motor neurons and iPSC neurons. Bulk RNA-sequencing and transcriptomic analysis of iPSC neurons within the scaffolds demonstrated coordinated upregulation of actin-remodelling, cell-matrix adhesion and metabolic pathways, indicative of a cytoskeletally adaptable neuron. When employed in an *ex vivo* dorsal root ganglia model, scaffold-mediated miR-133a inhibition promoted neurite extension and integration of delivered iPSC neurons with injured neural tissue. Finally, miR-133a-inhibitor-activated scaffolds upregulated neurovascular genes, increased endothelial cell migration and enhanced blood vessel formation *in vivo* in a chick embryo assay. These findings identify miR-133a as a neurotrophic target, elucidate the underlying mechanisms of action through transcriptomic analysis and demonstrate that biomimetic scaffold-mediated inhibition of miR-133a can enhance neuronal delivery for multifaceted spinal cord repair applications.

**Graphical Abstract:** 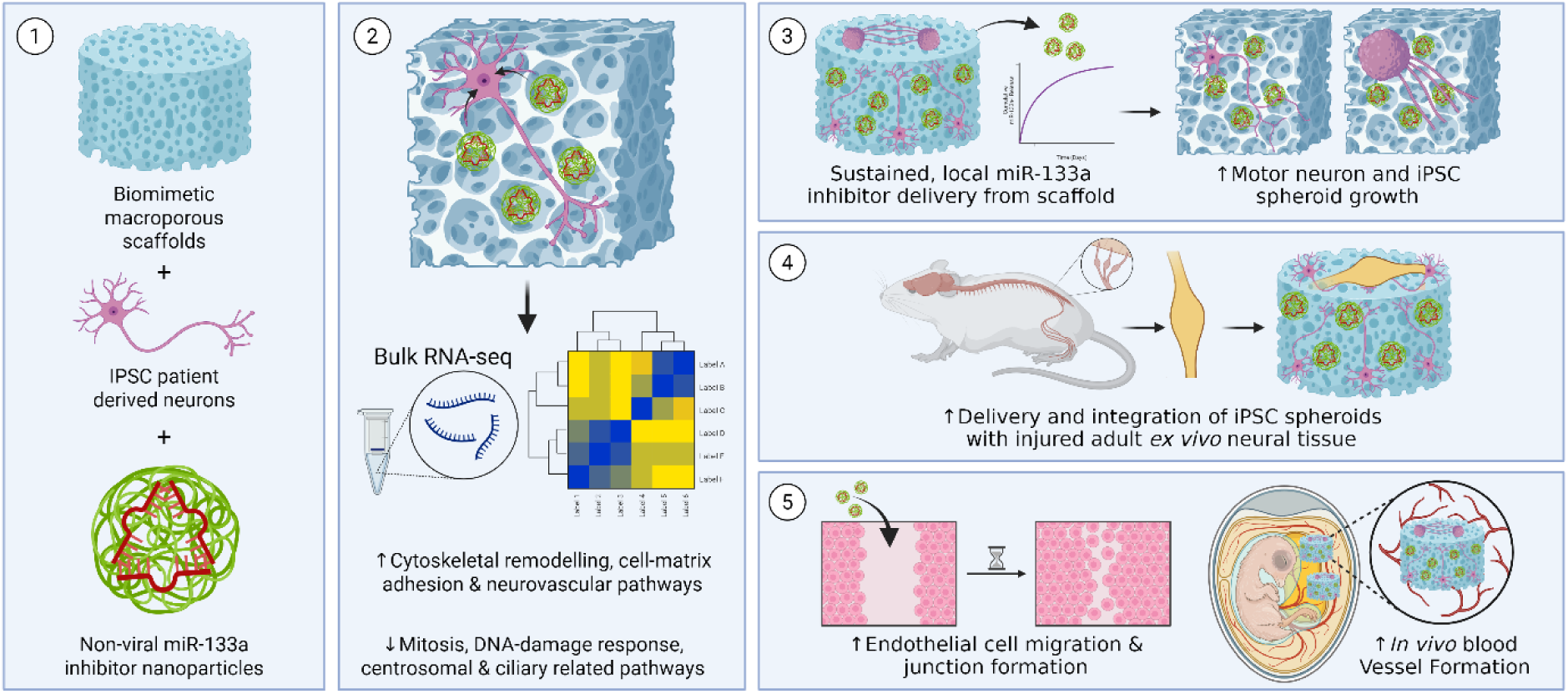

## Introduction

Spinal cord injury (SCI) is a traumatic injury where a lesion cavity develops, host neurons are lost and spared injured neurons cannot regrow[1]. These neurons cannot regrow their damaged axons in part due to the intrinsic capacity of adult neurons to inhibit actin mobilisation, ultimately attenuating neurite outgrowth[2,3]. Additionally, the inhibitory environment created by injury-responsive glial cells, along with the extensive loss of tissue suffered and the formation of a physical cavity create extrinsic barriers to neuronal regrowth[4–6]. Implantable biomaterial-based scaffold platforms have the potential to support localised, sustained delivery of therapeutic gene- and neural cell-cargo at the site of injury while physically bridging the injury site with a neurotrophic substrate[7]. Scaffold-based delivery of therapeutics and/or stem cells can promote host neuronal growth, replace lost cells and locally deliver neurons that can form functional relays across the injury site[1,8,9]. By designing biomimetic scaffold implants that mimic native spinal cord properties, key physicochemical cues such as matrix composition, stiffness and topography can be harnessed to stimulate neural tissue repair. Our previous work has developed soft, macroporous hyaluronic acid (HyA) scaffolds functionalised with collagen-IV (CIV) and fibronectin (FN) that reproduce the matrix composition and mechanics of healthy spinal cord and promote neurite extension[10] and immunomodulatory responses from astrocytes[11], providing a platform for spinal cord repair. However, while biomimetic implantable scaffolds can help overcome the extrinsic barriers of the unsupportive environs of the injured spinal cord and locally deliver cargo, there is still a need to boost the survival and effectiveness of delivered neurons to improve their integration with spared host tissue[1,9].

Combinatorial strategies have been widely employed to overcome the multifaceted challenges of spinal cord injury repair (i.e. using electrostimulation[12–14] and rehabilitation[15]) in parallel with the implantation of scaffolds and/or stem cell delivery[1,7]. The growing area of gene delivery to boost the effectiveness of delivered cells and remove intrinsic barriers to host neuronal growth has shown promise, especially when combined with scaffold platforms that enable localised, sustained release[16]. Many approaches to date have delivered genes to upregulate growth factor-related signals[17,18] or conversely inhibit intrinsic inhibitors restricting neurite outgrowth[19,20]. As a result, there is growing interest in the delivery of nucleic acids that can modulate multiple targets in neurons simultaneously. MicroRNAs (miRNAs), short non-coding RNA molecules which regulate post-transcriptional gene expression in cells, are an attractive cargo to target multiple genes and pathways within neurons[21]. Delivery of synthetic miRNAs (i.e. miRNA mimics or inhibitors) can upregulate or inhibit gene function by altering the expression of cassettes of genes. Unlike the direct delivery of growth factors or pharmaceutical inhibitors, miRNAs can be delivered in more physiologically relevant dosages and, by acting on multiple transcripts, can orchestrate changes in cytoskeletal and reparative programs rather than acting on a single gene[22].

Several miRNAs have been found to play key roles in regulating neural development and function, affecting neuronal maturation and neurite morphology[23]. One miRNA of particular interest is the miR-133 family (which encompasses miR-133a-1, miR-133a-2 and miR-133b), which when inhibited, has been shown to safeguard nerve cells from pharmacological insult through enhanced growth factor recognition[24], improved neural cell viability[25], and reduced neuropathic pain in rat nerves[26]. Additionally, miR-133a expression has been shown to affect cytoskeletal signalling in other peripheral cells[27], while targeting miR-133b in neurons enhances outgrowth by silencing neuronal actin-regulating genes[2,25]. Therefore, miR-133a inhibitor delivery might have potential to target actin regulation in neurons to enhance the survival, neurite extension and integration of transplantable induced pluripotent stem cell (iPSC)-derived neurons within supportive scaffold environments for spinal cord repair[16].

Despite the promise of miRNA-based therapies for neural repair, challenges remain in protecting miRNA from degradation while allowing for localised and sustained release to the site of interest. Our group has utilised gene-activated scaffold platforms for different tissue applications using non-viral cell-penetrating peptides to safely deliver gene cargos[28–30]. These gene-activated scaffolds leverage the bioactive environment of scaffolds and the protective role of non-viral vectors to deliver therapeutics at the site of clinical interest, reducing the need for large doses, overcoming systemic delivery challenges[29]. In this context, non-viral RALA cell-penetrating peptides[29,31] offer a versatile platform for complexing miRNA inhibitor cargo, enabling high loading efficiency and sustained release when incorporated into 3D scaffolds[29]. We hypothesised that by utilising RALA, miRNA-activated scaffolds could be manufactured with the potential to offer local delivery of therapeutic miR-133a inhibitor cargos to seeded neurons and injured neural cells in the surrounding environment[16], with the ability to modulate cytoskeletal pathways and influence processes such as angiogenesis which are required for neuronal survival.

Therefore, this study aimed to develop a biomimetic, macroporous, HyA-based scaffold platform that: (i) supports the growth of iPSC-derived neurons; (ii) enables RALA-mediated delivery and sustained release of miR-133a inhibitors and; (iii) enhances the growth and integration of iPSC neurons into injured neural tissue while modulating broader neuro-regenerative and angiogenic responses. By combining biomimetic scaffold design with miRNA-based gene activation and defining the underlying mechanism of action via transcriptomic profiling, we sought to develop and mechanistically characterise a multifaceted platform for next-generation spinal cord repair applications.

## Results

### Soft, biomimetic collagen-IV and fibronectin hyaluronic acid scaffolds promote iPSC neuronal growth

First, to confirm the optimal scaffold matrix composition for promoting neuronal outgrowth, NSC-34 mouse motor neuron-like cells were cultured on poly-L-lysine (PLL, an inert substrate) and a combination of collagen-IV (CIV) and fibronectin (FN) (previously shown to promote neural cell growth individually at both whole cell and nanoscale levels[11,32]) substrates and their morphology was assessed (Figure 1A). Neurons grown on CIV/FN substrates showed elongated morphologies with long neurites extending compared to PLL-grown neurons, which were rounded and showed little neurite extension (Figure 1B-C). When changes in neuronal morphology were quantified, both the average and max neurite length were significantly enhanced when grown on CIV/FN compared to PLL by ∼106% and ∼197%, respectively (Figure 1D-E). Qualitatively, nanoscale changes in actin protrusion were also observed in neurons grown on CIV/FN, where sub-micron protrusions extended, while neurons on PLL demonstrated fewer actin protrusions in comparison (Figure 1F).

**Figure 1.**
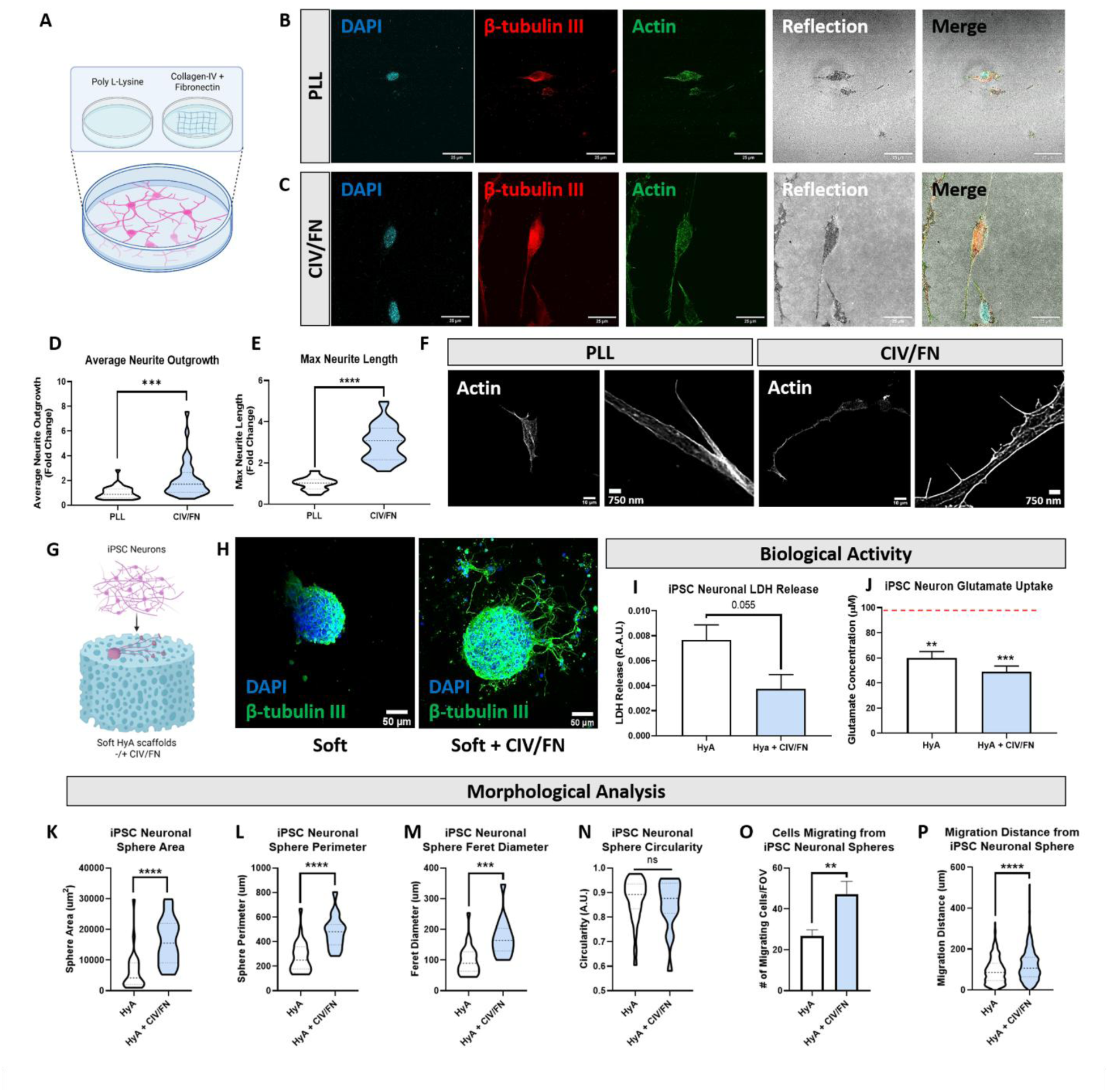
Soft, collagen-IV & fibronectin scaffolds enhance iPSC neuronal growth. A) Neurons were grown on poly-L-lysine (PLL) and collagen-IV/fibronectin (CIV/FN) coated substrates. B-C) Confocal and reflection imaging of neurons grown on different substrates, on PLL and CIV/FN substrates, reveals distinct morphological differences. Scale bar = 25 µm. D-E) CIV/FN substrates significantly enhanced neuronal max and average neurite length compared to neurons grown on PLL only. F) Nanoscale protrusions in neuronal actin extension are enhanced on CIV/FN substrates. Scale bars = 25 µm & 750 nm G) Experimental outline of iPSC neurons seeded in soft hyaluronic acid (HyA) scaffolds with and without CIV/FN functionalization. H) HyA, CIV/FN scaffolds enhance neurite extension from iPSC neuronal spheroids. Scale bar = 50 µm. I) Cytotoxic LDH release from iPSC neurons is reduced in HyA, CIV/FN scaffolds compared to HyA-only scaffolds. J) HyA, CIV/FN scaffolds significantly enhanced the ability of iPSC neurons to uptake glutamate more than neurons in HyA-only scaffolds compared to baseline values. K-M) Hya, CIV/FN scaffolds significantly enhanced iPSC neuronal sphere area, perimeter and feret diameter (elongation) compared to HyA only scaffolds. N) Neuronal spheroid area was not affected by scaffold composition. O-P) HyA, CIV/FN scaffolds significantly enhanced the number and distance of migrating cells from spheroids compared to HyA scaffolds. All data N=3. Analysis via unpaired two-tailed t-tests.

Once, CIV/FN were confirmed to promote neuronal outgrowth, soft, hyaluronic acid (HyA) macroporous scaffolds that mimic the stiffness, matrix composition and topography of the native spinal cord tissue[10,11,33] were functionalized with CIV/FN. Next, human iPSC-derived neurons were seeded on these HyA scaffolds with and without CIV/FN functionalization (Figure 1G). After 7 days of scaffold culture, iPSC neurons were found to form spheroids that extended several long neurites over time in soft, CIV/FN scaffolds whereas scaffolds without CIV/FN did not extend as many long neurites (Figure 1H). Following analysis of the biological activity of the iPSC neurons grown in the scaffolds, it was found that iPSC neurons in HyA, CIV/FN scaffolds showed a trend towards lower levels of cytotoxic LDH release (Figure 1I). When looking at the capacity of iPSC neurons to uptake glutamate (a key functional process of neurons), it was found that iPSC neurons had higher glutamate uptake in HyA, CIV/FN scaffold groups compared to HyA scaffolds only (Figure 1J). When the morphology of the iPSC neurons was analysed, spheres with significantly larger area, perimeter and feret diameter (the longest distance between any two points) were observed in HyA, CIV/FN functionalized scaffolds while maintaining a similarly rounded spheroid structure (Figure 1K-N). Additionally, the migration of cells from the iPSC neuronal spheroids was also significantly enhanced in HyA, CIV/FN groups, where the number of cells that migrated and the max distance of migration was significantly enhanced in HyA, CIV/FN scaffolds (Figure 1O-P). Overall, HyA, CIV/FN scaffolds supported iPSC neuronal growth, neurite extension and migration from scaffold-formed spheroids.

### Bioinformatic in silico identification of miR-133a as a neuron-related cytoskeletal target and characterisation of miR-133a inhibitor nanoparticles and scaffold release kinetics

Once the optimal scaffold physicochemical properties to support iPSC neuronal growth were established, we next sought to confirm whether inhibition of miR-133a impacted neuronal actin dynamics and neurite growth. Pathway enrichment analysis was performed by compiling data from different published public databases into DAVID software to analyse significantly enriched pathways in which miR-133a is involved (Figure 2A). First, gene-ontology analysis found that miR-133a affects several key neuronal-specific pathways involved in neurotransmitter transport, neuronal cell body organisation, metabolism, GABA-ergic synapse formation and actin filament organisation (Figure 2B). Through performing Kyoto encyclopaedia of genes and genomes (KEGG) pathway analysis on the collected datasets, synaptic vesicle cycling, regulation of the actin cytoskeleton and the PI3K-AKT signalling pathway were also observed to be affected by miR-133a (Figure 2C & Supplemental Figures 1-3).

**Figure 2.**
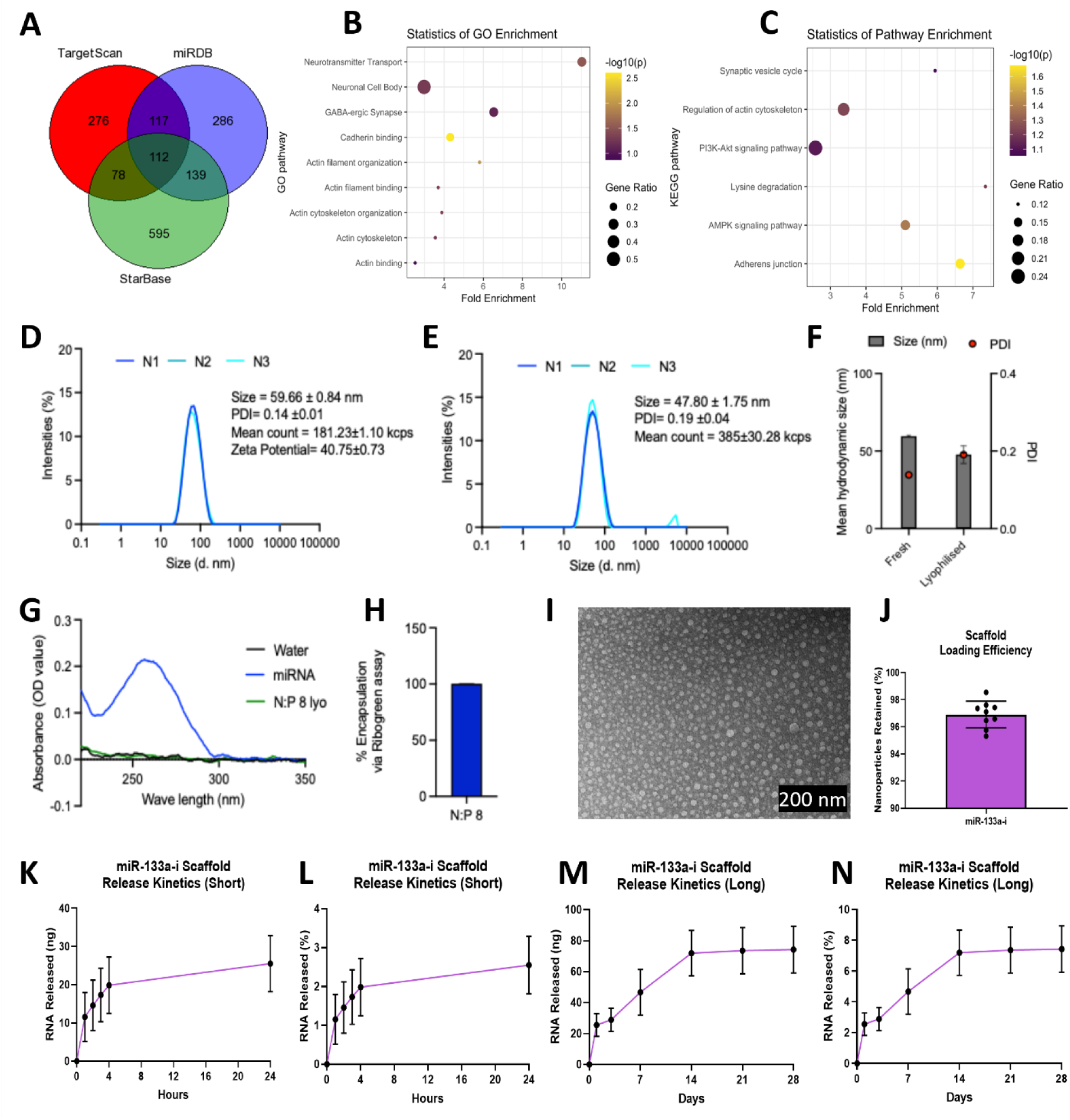
Bioinformatic mapping of miR-133a neuronal pathways and physicochemical characterisation of miR-133a inhibitor nanoparticles and scaffold release kinetics. A) Venn diagram of predicted miR-133a targets identified across three public miRNA–mRNA interaction databases (TargetScan, miRDB and StarBase), showing the shared gene set used for downstream enrichment analysis. B) Gene Ontology (GO) enrichment analysis of common predicted miR-133a targets, highlighting over-represented neuronal and cytoskeletal categories. C) KEGG pathway enrichment analysis of the same gene set, demonstrating enrichment of pathways linked to synaptic vesicle cycling, regulation of the actin cytoskeleton and adherens junctions. D-E) Size distribution spectra of fresh and lyophilised RALA/miR-133a-i nanoparticles formulated at an N:P 8 ratio. Insert summarises z-average, polydispersity index (PDI), mean count rate and zeta potential +/- SEM. F) Mean hydrodynamic size and PDI of fresh and lyophilised RALA/miR-133a-i nanoparticles at N:P ratio of 8. G) Ion exchange chromatography results showing absorbance spectra of RALA/ miR-133a-i nanoparticles, water and unencapsulated miRNA following chromatography through an anionic Sephadex resin. H) Encapsulation efficiency of RALA/miR-133a-i nanoparticles measured by quantifying free miRNA present in solution. I) TEM image showing RALA/miR-133a-i nanoparticles. J) Loading efficiency of >96% of nanoparticles in hyaluronic acid scaffolds was detected by the Ribogreen Assay. Nanoparticle release kinetics measured in ng and percentage change for short-term release (K-L) and long-term release periods (M-N) demonstrated an initial burst release followed by a slower, consistent release.

As miR-133a was found to be associated with key neuronal actin-regulating and growth-promoting pathways, we next sought to formulate non-viral miR-133a inhibitor (miR-133a-i) nanoparticles that could be delivered to neurons and repress miR-133a expression. To encapsulate the miR-133a-i cargo for delivery, the miR-133a-i was complexed with the cell-penetrating peptide RALA[29,31]. Characterisation of fresh and lyophilised nanoparticles indicated an optimal size (∼47.8-59.7 nm), polydispersity index (0.14-0.19) and zeta potential (∼40.75 mV) for cell internalisation (Figure 2D-E). An optimal N:P ratio of 8[29] was also found to provide an encapsulation efficiency of >95% (Figure 2F-G). Transmission electron microscopy imaging also revealed a uniform size distribution of the produced miR-133a-i nanoparticles (Figure 2H). Next, the developed miR-133a-i nanoparticles were seeded into the soft HyA, CIV/FN scaffolds (with stiffness of <1.5kPa as established in previous studies[10,11,13,33]) to produce the miR-133a-i-activated scaffolds with a loading efficiency of ∼96.9% of the nanoparticles (Figure 2I). As the sustained and controlled release of the nanoparticles from the scaffold is key for successful transfection, we next looked at the short and long-term release profiles up to 28 days *in vitro*. The scaffolds released ∼25.5 ng of the 1 µg of loaded RNA within the first 24 hours (Figure 2J), which corresponded to a cumulative release of ∼2.55% of the total RNA cargo (Figure 2K). The initial release phase was then followed by a continued release of nanoparticles up to 14 days, which plateaued between days 14-28 with a final release of ∼74.3 ng of RNA of 7.43% of the total loaded RNA (Figure 2L-M). Overall, the results show that the HyA, CIV/FN scaffolds effectively incorporated the miR-133a-i nanoparticles, providing localised and sustained delivery up to 28 days.

### Biomimetic miR-133a-i-activated scaffolds provide localised delivery to motor neurons and enhance neurite outgrowth

Having confirmed the effective incorporation and controlled release of the miR-133a-i nanoparticles from the biomimetic scaffolds, we next aimed to determine whether these miR-133a-i-activated scaffolds successfully promote changes in neurite outgrowth without compromising neuronal biocompatibility (Figure 3). Scanning electron microscopy revealed that motor neuron-like cells within non-activated nanoparticle-free scaffolds had smooth cell surfaces with rounded cell morphologies (Figure 3A). In contrast, neurons seeded within miR-133a-i-activated scaffolds were found to co-localise with nanoparticles that decorated the cell surfaces (Figure 3B). Confocal microscopy revealed differences in neuronal coverage of the scaffolds, where the miR-133a-i-activated scaffolds promoted larger areas of neuronal coverage and cell numbers (Figure 3C). Analysis of neuronal metabolic activity revealed that miR-133a-i-activated scaffolds significantly enhanced neuronal metabolic activity and LDH release (Figure 3D-E). Following morphological analysis, it was observed that the neuronal area coverage was significantly enhanced in miR-133a-i-activated scaffolds compared to untreated scaffolds (Figure 3F). Finally, analysis of neuronal outgrowth via measuring the max neurite length demonstrated that the miR-133a-i-activated scaffolds significantly enhanced neurite outgrowth compared to untreated scaffold groups (Figure 3G). Overall, biomimetic soft HyA, CIV/FN scaffolds supported the local delivery of nanoparticles to neurons and a combination of optimal scaffold properties and miR-133a-i delivery enhanced neurite outgrowth.

**Figure 3.**
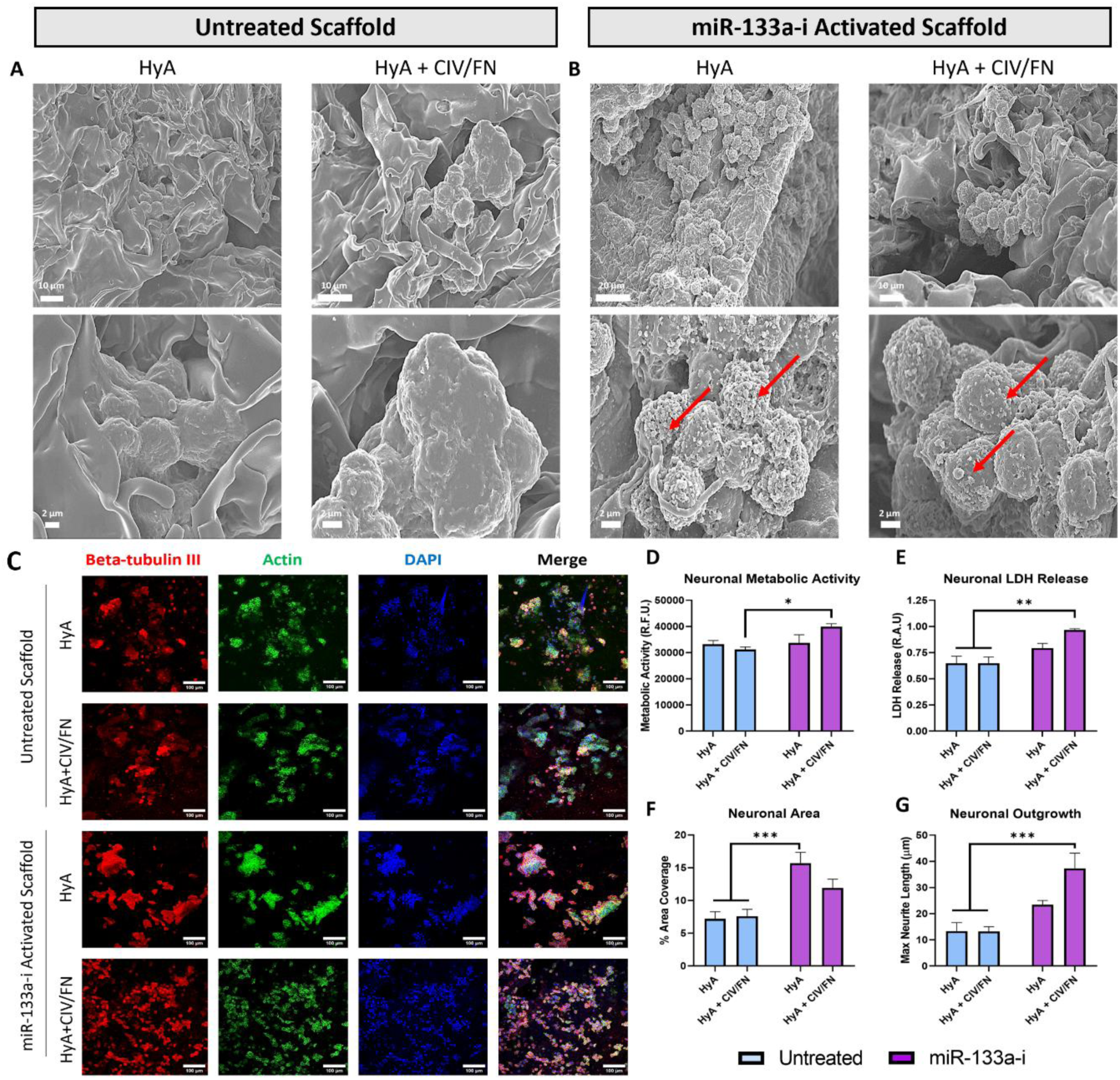
Biomimetic miR-133a-i-activated scaffolds facilitate local nanoparticle delivery to enhance neurite outgrowth. A-B) Scanning electron microscopy of mouse motor neuron-like cells in untreated scaffolds shows smooth cell surfaces, while cells in miR-133a-i-activated scaffolds co-localise with nanoparticles (red arrows indicate nanoparticles). A&B) Scale bars = 10 & 20 µm top row respectively, 2 µm bottom row for all. C) Confocal microscopy of neurons shows larger areas of scaffold coverage in miR-133a-i-activated scaffolds compared to untreated scaffolds. Scale bars = 100 µm. D-E) Neuronal metabolic activity and LDH release are significantly enhanced in miR-133a-i-activated HyA scaffolds functionalized with CIV/FN compared to untreated scaffolds. F) Neuronal area coverage is significantly increased in miR-133a-i-activated scaffolds compared to untreated scaffolds. G) Max neurite length is significantly increased in miR-133a-i-activated HyA scaffolds functionalized with CIV/FN compared to untreated scaffolds. All analyses via two-way ANOVA, Bonferroni post-hoc test. N=3 for all data.

### Biomimetic miR-133a-i-activated scaffolds provide localised delivery to and enhance iPSC neuronal spheroid size and outgrowth

Once it was established that miR-133a-i-activated scaffolds functionalized with CIV/FN enhanced neurite outgrowth, iPSC neurons were seeded within the untreated and gene-activated scaffolds to determine if scaffold-mediated miR-133a inhibition enhanced their outgrowth for delivery to neural tissue. First, scanning electron microscopy revealed that iPSC neurons formed spheroids within both untreated and miR-133a-i-activated scaffolds, where spheroids in miR-133a-i-activated scaffolds were larger in size and co-localised with nanoparticles (Figure 4A-B). When the morphology of the iPSC neurons within the scaffolds was analysed via confocal microscopy, distinct morphological differences were observed where spheroids in miR-133a-i-activated scaffolds showed higher levels of neurite extension compared to neurons in untreated scaffolds, which showed rounded spheroid morphologies with few protrusions (Figure 4C-D). Analysis of iPSC neuronal LDH release and metabolic activity showed no significant difference between scaffold groups, indicating that the miR-133a-i-activated scaffolds did not affect viability (Figure 4E-F). Analysis of morphology revealed significant increases in iPSC neuronal spheroid size, perimeter, elongation and neurite extension compared to the untreated scaffolds (Figure 4G-J).

**Figure 4.**
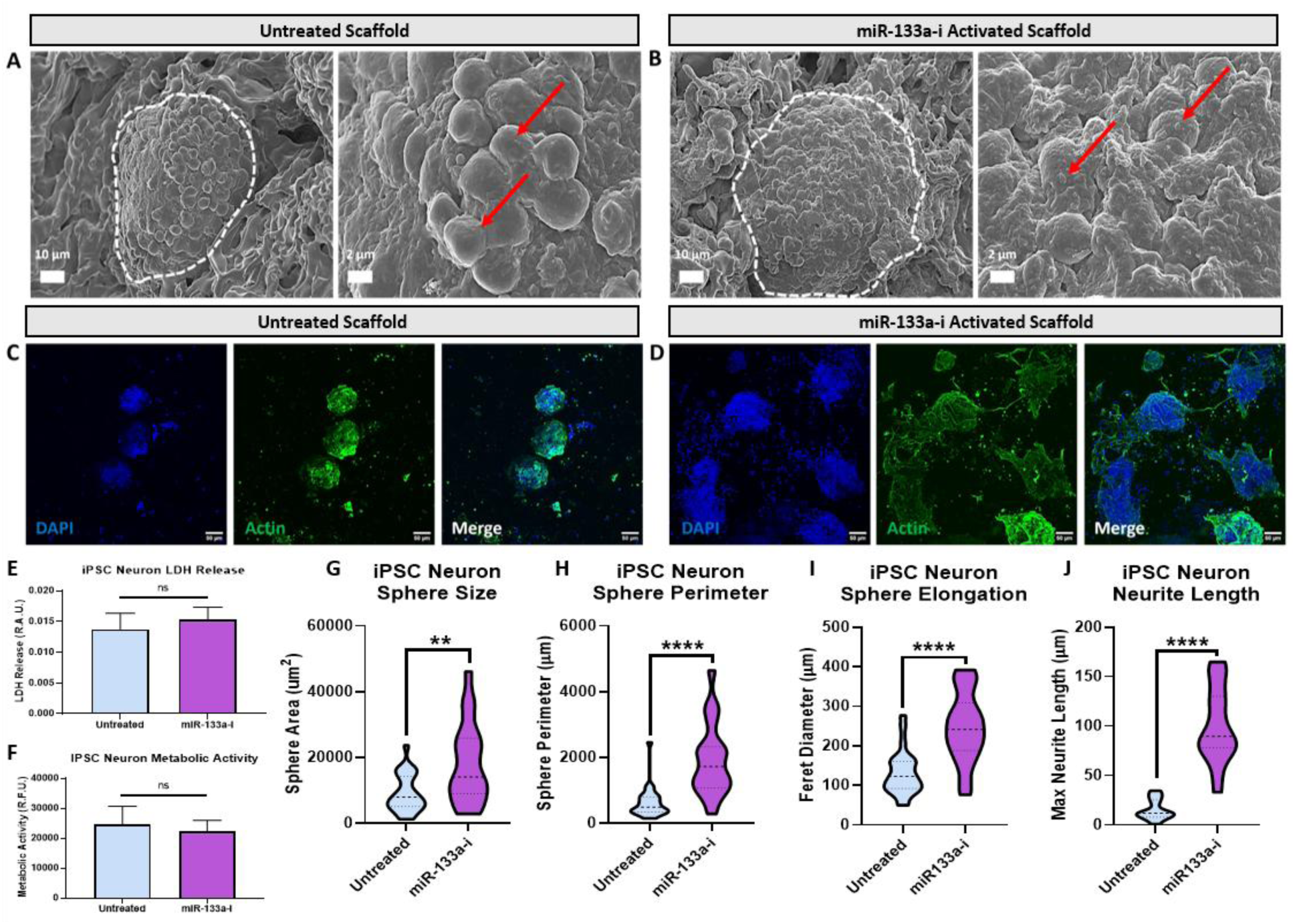
Biomimetic miR-133a-i-activated scaffolds enhance iPSC neuronal spreading and neurite extension. A-B) Scanning electron microscopy reveals IPSC neurons with smooth surfaces within condensed spheroids in untreated scaffolds, while iPSC neurons in miR-133a-i-activated scaffold neuronal spheroids are observed to spread, and individual neurons co-localise with nanoparticles. Scale bars = 10 µm & 2 µm. C-D) Confocal microscopy reveals that iPSC neuronal spheroids are larger and extend neurites between each other in miR-133a-i-activated scaffolds compared to untreated scaffolds. C-D) Scale bars = 50 µm. E-F) Metabolic activity and cytotoxicity of iPSC neurons are not affected in miR-133a-i-activated scaffolds. G-J) iPSC neuronal spheroid size, perimeter, elongation and neurite extension are significantly enhanced in miR-133a-i-activated scaffolds compared to untreated scaffolds. All data N=3. All analyses via an unpaired two-tailed t-test.

### Biomimetic miR-133a-i-activated scaffolds promote neuronal actin dynamics, cell-matrix adhesion and endocytosis-related gene expression

Having confirmed that the developed miR-133a-i-activated scaffolds successfully promoted neurite outgrowth in both mouse motor neurons and iPSC-derived neurons, we next aimed to define the underlying genetic mechanisms driving the changes in neuronal morphology. Transcriptomic profiling of iPSC-derived neurons using bulk RNA-sequencing following scaffold-mediated miR-133a inhibition revealed a coordinated shift in gene expression related to neurite outgrowth and structural remodelling. First, a significant decrease in miR-133a expression was observed in iPSC neurons in miR-133a-i-activated scaffolds compared to iPSC neurons in untreated scaffolds (Figure 5A). When analyzing largescale genomic changes, 1282 differentially enriched genes were identified following scaffold-mediated miR-133a inhibition (703 upregulated, 573 downregulated) (Figure 5B-C).

**Figure 5.**
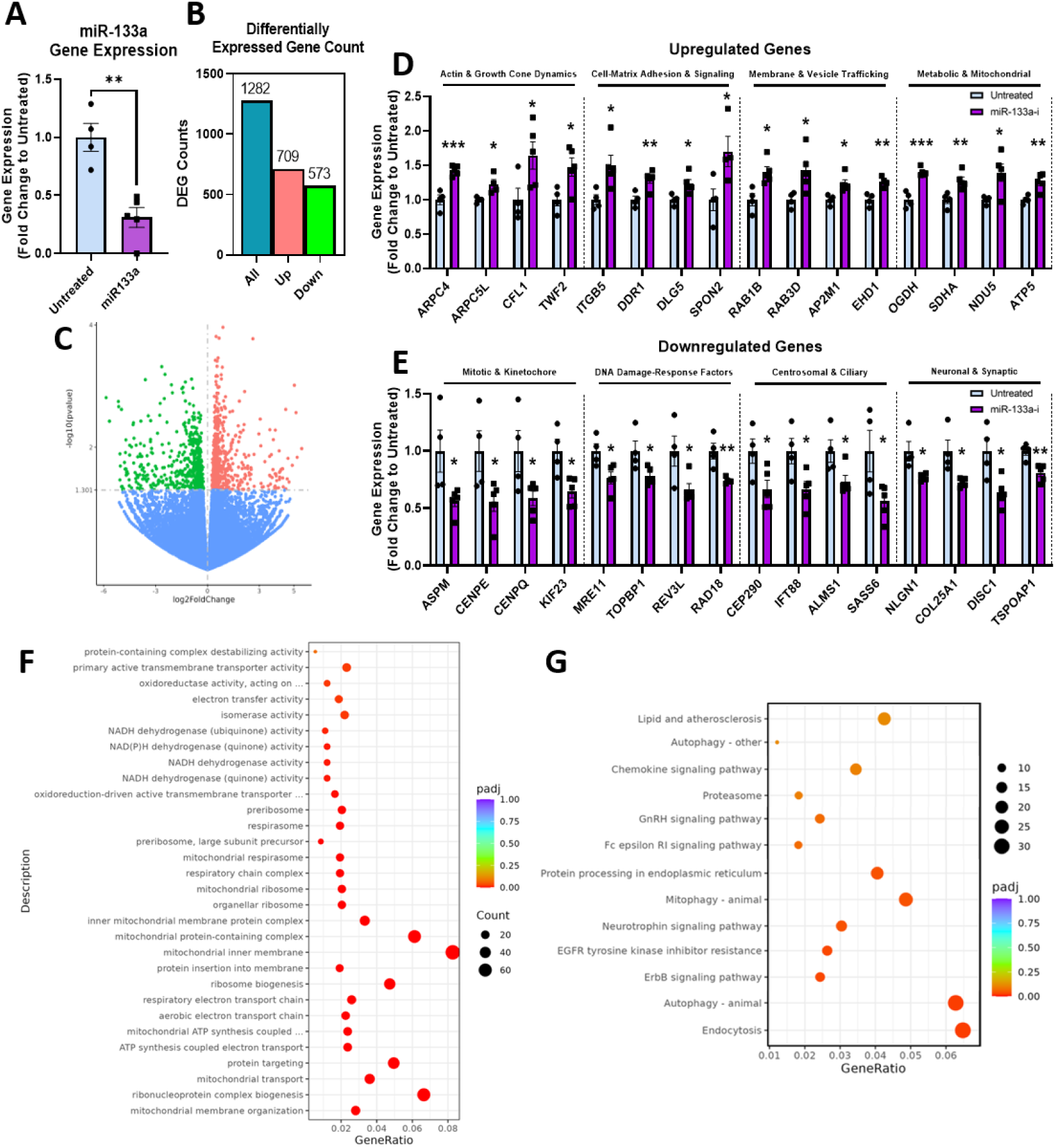
Transcriptomic profiling of iPSC neurons in miR-133a-i-activated scaffolds reveals modulation of actin dynamics, matrix signalling, metabolic and endocytosis pathways. A) MiR-133a gene expression levels are significantly reduced in iPSC-derived neurons within miR-133a-i-activated scaffolds compared with untreated scaffolds. B) Number of differentially expressed iPSC neuronal genes in miR-133a-i-activated scaffold groups compared to untreated scaffolds. C) Volcano plot of bulk RNA-sequencing data comparing differentially expressed genes in iPSC neurons in miR-133a-i-activated versus untreated scaffolds, highlighting significantly up-regulated (red) and down-regulated (green) genes (adjusted p-value threshold indicated by horizontal line). D) Normalised iPSC neuronal gene expression changes for representative significantly upregulated genes involved in actin and growth cone dynamics, cell matrix & adhesion signalling, membrane & vesicle trafficking and neuronal metabolic-related genes in untreated versus miR-133a-i-activated scaffolds. E) Normalised gene expression changes for representative significantly downregulated genes involved in mitosis, DNA-damage response, centrosomal & ciliary dynamics and neuronal & synaptic related genes in untreated versus miR-133a-i-activated scaffolds. F) Gene Ontology (GO) enrichment analysis of significantly differentially expressed genes, demonstrating over-representation of terms related to mitochondrial function, protein translation and cellular metabolic activity. G) KEGG pathway enrichment analysis of the same gene set, indicating enrichment of pathways associated with neurotrophin signalling, autophagy, endocytosis and other neuroregenerative or stress-response pathways in neurons within miR-133a-i-activated scaffolds. Differential expression and enrichment analyses were performed on bulk RNA-seq datasets (N=4-5). A, D-E) Analysis for each gene performed via unpaired, two-tailed t-tests where *p<0.05, **P<0.01 & ***p<0.001.

Genes involved in actin dynamics and growth-cone motility (e.g. *ARPC4, ARPC5L*), cell–matrix adhesion and signalling (e.g. *ITGB5, DDR1*), and membrane/vesicle trafficking (e.g. *RAB1B, RAB3D*) were upregulated in miR-133a-i-activated scaffolds, alongside broad enhancement of mitochondrial and metabolic pathways (e.g. *OGDH, SDHA*), suggesting increased energetic support for neurite extension (Figure 5D). In parallel, several kinases and transcriptional regulators previously linked to neuritogenesis and structural plasticity (e.g. *CDK5, BCAR1*) were also induced (Supplemental Figure 4). Together, these changes possibly define a gene-expression program that facilitates enhanced neurite growth and remodelling in response to scaffold-mediated miR-133a inhibition in patient-derived iPSC neurons. In contrast, cell-cycle, DNA repair, cilia/centrosome and neuronal synaptic-associated genes were downregulated, consistent with a shift toward a more post-mitotic, projection-focused neuronal state elicited by miR-133a-i-activated scaffolds (Figure 5E). Together, these changes define a genotypic program that facilitates neurite growth and remodelling in response to the miR-133a-i-activated scaffolds.

To obtain an overview of the biological processes influenced by the miR-133a-i-activated scaffolds, we performed over-representation analysis on the significantly enriched genes (padj<0.05). Gene Ontology terms enriched in this set were dominated by mitochondrial and respiratory chain functions, including oxidoreductase and NADH dehydrogenase activity, components of the electron transport chain, mitochondrial inner membrane and respiratory chain complexes, as well as ATP synthase and multiple ribosomal and ribosome-biogenesis terms (Figure 5F). These data indicate that miR-133a-i-activated scaffolds drive a broad upregulation of mitochondrial oxidative phosphorylation, ATP production and translational capacity in iPSC-derived neurons. Consistent with this, KEGG pathway analysis of enriched genes (Figure 5G) highlighted enrichment of proteostasis and organelle-quality-control pathways such as the proteasome, autophagy and mitophagy, together with signalling and trafficking pathways including endocytosis, ErbB and EGFR-related signalling and neurotrophin signalling. Additional enrichment in lipid and atherosclerosis-related pathways also suggests coordinated remodelling of metabolic programs. These analyses show that miR-133a-i-activated scaffolds promote a pro-metabolic, pro-proteostatic and cytoskeletally plastic and signalling-competent state that is likely to support enhanced neurite growth and structural remodelling, addressing key intrinsic barriers to axonal regeneration in SCI repair.

### Biomimetic, miR-133a-i-activated scaffolds enhance iPSC neuronal delivery to injured neural tissue

Once we established a system-level effect of the miR-133a-i-activated scaffolds on iPSC neurons and demonstrated that these scaffolds promote neurite outgrowth, we next sought to determine whether the biomimetic miR-133a-i-activated scaffolds enhance iPSC neuronal delivery to injured tissue. Adult mouse dorsal root ganglia (DRG, an *ex vivo* model of injured neurite outgrowth) were cultured on the iPSC neuronal-seeded scaffolds with and without miR-133a-i activation (Figure 6A). Analysis of LDH release and pro-inflammatory tumour necrosis factor alpha (TNFα) and interleukin-1 beta (IL-1β) and anti-inflammatory interleukin-10 (IL-10) release showed no significant differences across the different scaffold groups, indicating miR-133a-i delivery did not exacerbate neural cytotoxicity or inflammatory responses (Figure 6B-E). Additionally, no differences in the uptake of glutamate compared to the baseline media levels were observed between the scaffold groups, indicating no major changes in glycolytic metabolic function (Figure 6F). Imaging via scanning electron microscopy showed that DRGs attached and integrate within the scaffold architecture (Figure 6G). Similarly, iPSC neurons were observed to form spheroids within the same scaffold groups (Figure 6H). Imaging of individual neural cells showed that cells formed uniform monolayers colonising the scaffold surface and that neural cells within miR-133a-i-activated scaffolds co-localised with nanoparticles (Figure 6I). Furthermore, long, dense neurites were observed extending across the surface of miR-133a-i-activated scaffolds, where untreated scaffolds did not extend such neurites (Figure 6J). Finally, confocal microscopy revealed that iPSC neurons in miR-133a-i-activated scaffolds were delivered to DRGs and extended neurites between the spheroids and DRG tissue, where untreated scaffolds did not (Figure 6K). Overall, the miR-133a-i-activated scaffolds successfully enhanced the delivery of iPSC neurons to injured neural tissue, promoting neurite extension and tissue integration.

**Figure 6.**
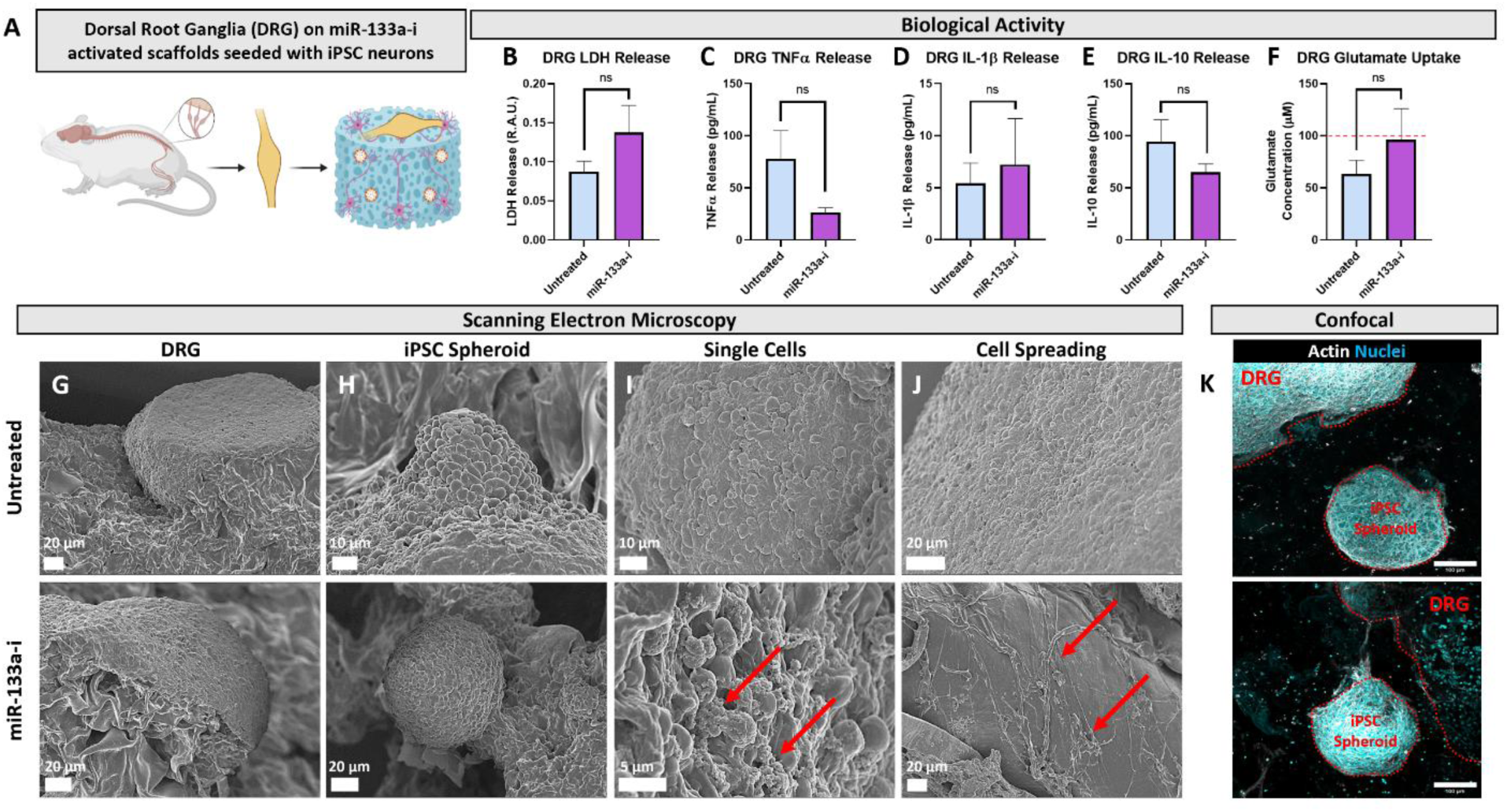
Biomimetic miR-133a-i-activated scaffolds enhance iPSC neuronal delivery to injured neural tissue. A) Experimental outline of dorsal root ganglia (DRG) culture on iPSC neuronal-seeded scaffolds. B-E) Analysis of cytotoxic LDH release, pro-inflammatory TNFα and IL-1β and anti-inflammatory IL-10 cytokine release showed no differences between untreated and miR-133a-i-activated scaffold groups. G-J) Scanning electron microscopy images of (G) DRG bodies, (H) iPSC neuronal spheroids, (I) single migrating cells, (J) cell spreading and neurite extensions within untreated and miR-133a-i-activated scaffolds. K) Confocal microscopy reveals enhanced neuronal extension between DRGs and iPSC neurons within miR-133a-i-activated scaffolds, while no extension was observed in untreated scaffolds. Scale bar = 100 µm. All data N=3. Analysis via unpaired two-tailed t-test.

### Neurovascular miR-133a inhibition augments endothelial migration and stimulates angiogenesis *in vivo*

Having shown that miR-133a-i-activated scaffolds enhanced iPSC neuronal engraftment and neurite integration within neural tissue, we next sought to assess whether this strategy modulated angiogenic processes, a key requirement for functional neural repair. To explore whether miR-133a inhibition engages angiogenesis-related transcriptional programs, we examined expression of vascular and adhesion-associated genes in miR-133a-i-activated scaffold grown iPSC neurons. Analysis of significantly regulated gene transcripts revealed robust upregulation of secreted and growth-factor-like angiogenic molecules, matrix adhesion and junctional regulators, and inflammatory/immune modulators linked to angiogenic responses, together with the neurovascular transporter MFSD2A and vascular barrier related genes (Figure 7A). These changes indicate that neuronal miR-133a-i-activated scaffolds may induce a pro-angiogenic gene expression signature indicative of a microenvironment for vascular remodelling.

**Figure 7.**
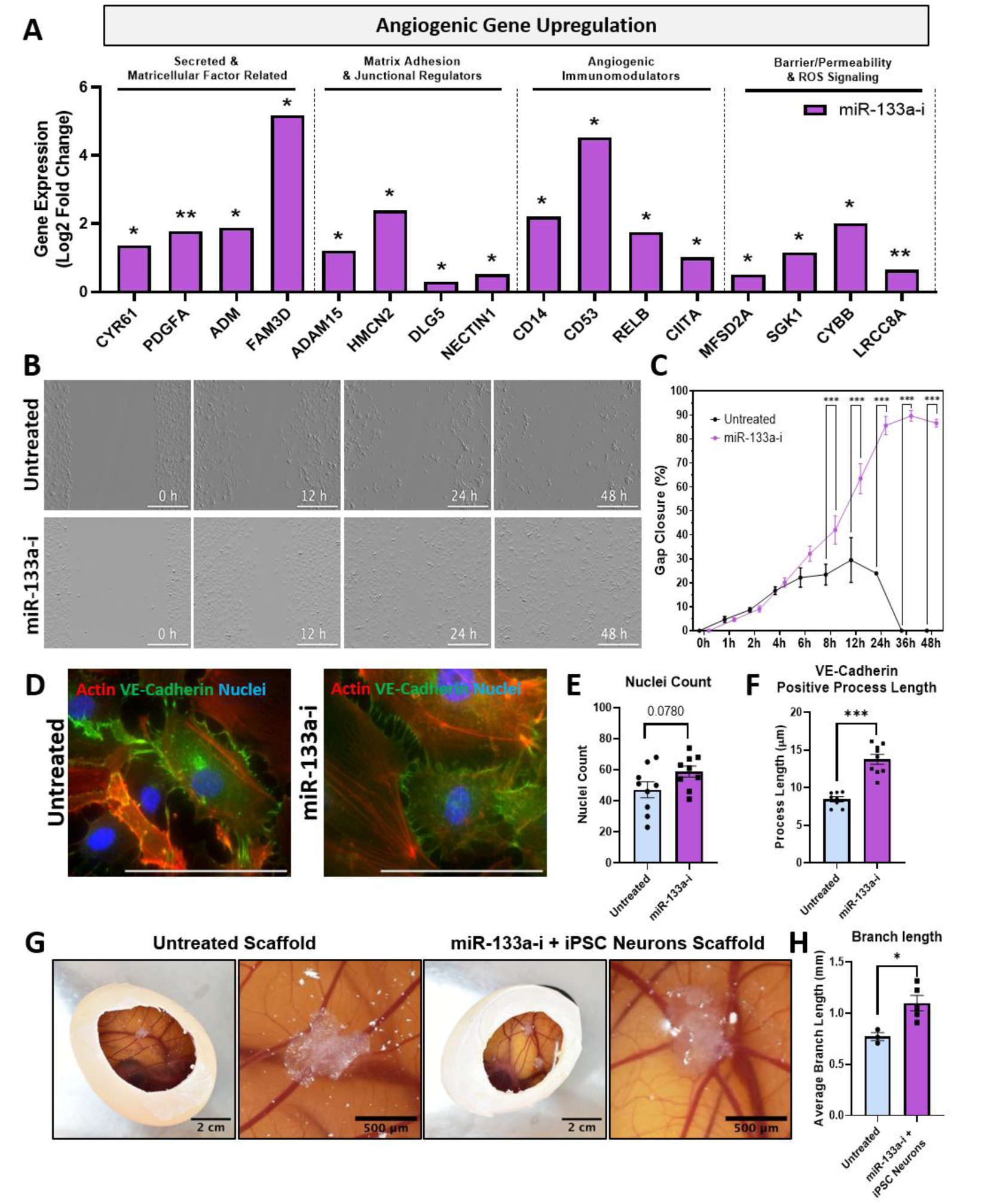
Biomimetic miR-133a-i-activated scaffolds promote neurovascular gene signatures, endothelial cell actin mobility and *in vivo* chick embryo blood vessel formation. A) Differentially expressed genes associated with angiogenesis and neurovascular signalling from bulk RNA-seq of iPSC neurons cultured within untreated versus miR-133a-i-activated scaffolds, demonstrating upregulation of pro-angiogenic and vessel-stabilising factor-related genes. B-C) Scratch assay of endothelial cell monolayers with representative live cell images from 0-48 h shows enhanced migration and scratch closure when treated with miR-133a-i cargo. Scale bar = 500 µm. D) Immunofluorescence images of endothelial cells stained for F-actin (red), VE-cadherin (green), and nuclei (blue) following treatment with miR-133a-I cargo. Scale bar = 100 µm. E) Nuclei counts and F) analysis of endothelial cell processes confirming increased cytoskeletal dynamics and junction formation in response to miR-133a-i cargo. G) Visualisation of *in vivo* chick embryo vasculature formation around implanted scaffolds. Scale bars = 2 cm & 500 µm. H) MiR-133a-i-activated scaffolds seeded with iPSC neurons significantly enhance vasculature branch length compared to untreated scaffolds. A) N=4-5, data presented as log2 fold change to untreated with adjusted p-(padj) values of differentially expressed genes. C) N=3, analysed via two-way ANOVA with a Bonferroni post-hoc test. E-F, H) Analysed via unpaired, two-tailed t-test. Data are presented as mean ± SEM; *p < 0.05, **p < 0.01, ***p < 0.001 where indicated.

We next assessed whether miR-133a-i delivery directly influences endothelial behaviour. Human umbilical vein endothelial cell (HUVEC) monolayers treated with miR-133a-i cargo showed markedly accelerated closure in a scratch-wound assay compared with untreated controls, indicative of enhanced collective migration (Figure 7B–C). Quantification of endothelial proliferation following miR-133a inhibition further demonstrated a trending increase in cell number over 48 h (Figure 7D-E). Immunofluorescence staining revealed more prominent VE-cadherin–positive junctions and pronounced actin stress fibres and intercellular extensions in miR-133a-i-treated cultures, consistent with the formation of continuous, elongated endothelial networks (Figure 7D & F). Together, these data show that miR-133a inhibition augments endothelial motility, growth and junctional organisation, features expected to support the formation of more extensive and potentially more stable vascular networks.

Finally, to determine whether the soft biomimetic iPSC neuron-seeded miR-133a-i-activated scaffold promote angiogenesis *in vivo*, we employed the chick chorioallantoic membrane (CAM) assay. Scaffolds functionalized with miR-133a-i nanoparticles and iPSC neurons induced a visibly denser and more radially organised vascular plexus around the implantation site compared with untreated, cell-free control scaffolds (Figure 7G). Quantitative analysis of vessel architecture demonstrated a significant increase in average branch length in the miR-133a-i-activated scaffold group, indicating enhanced sprouting and growth of blood vessels (Figure 7H). Collectively, these findings suggest that miR-133a-i-activated scaffolds not only support neuronal integration but also engage neurovascular programs that promote endothelial migration, junctional remodelling and angiogenesis *in vivo*.

## Discussion

The objective of this study was to develop a biomimetic miR-133a-i-activated scaffold to enhance the delivery and growth of iPSC-derived neurons for multifaceted repair of injured neural tissue. Building on our previous work developing soft, macroporous HyA scaffolds functionalised with collagen-IV and fibronectin, which provide a mechanically matched, neurotrophic environment, we here integrated the non-viral RALA cell-penetrating peptides complexed with miR-133a inhibitors to create a gene-activated platform with high loading efficiency and sustained release over 28 days. Bulk RNA sequencing and bioinformatic analyses showed that miR-133a-i-activated scaffolds drive iPSC-derived neurons towards a state enriched for actin cytoskeletal remodelling, cell–matrix adhesion, vesicle trafficking and neurotransmitter-related pathways, consistent with the observed enhancement of neurite outgrowth in both mouse motor neurons and iPSC neurons cultured within these scaffolds. When applied in an injured dorsal root ganglia model, the miR-133a-i-activated scaffolds enhanced iPSC neurite extension and structural integration with host neural tissue. Angiogenic assays further demonstrated that miR-133a inhibition promotes endothelial migration, actin mobilisation and *in vivo* blood-vessel formation, indicating that the miR-133a-i-activated scaffold microenvironment engages supportive neurovascular programmes. Together, we show the development of a biomimetic miR-133a-i-activated scaffolds as a platform that simultaneously targets intrinsic cytoskeletal barriers to axonal growth and vascular support, thereby augmenting iPSC neuronal integration into injured tissue for spinal cord injury repair applications.

The localised delivery of patient-derived neurons for neural repair applications has remained a challenge in part due to the harsh microenvironment of the injury site in spinal cord injury, resulting in low viability of the delivered cells[9,34]. Here, we show that soft, biomimetic macroporous hyaluronic acid scaffolds functionalized with neurotrophic ECM proteins, collagen-IV and fibronectin, can promote the growth of sensitive iPSC neurons within 3D environments for delivery. Other studies that have delivered iPSC-derived neurons to the injured spinal cord using scaffolds that mimic the native spinal cord tissue properties have demonstrated successful integration of iPSC neurons and improvements in function following transplantation in rodents[34]. For example, Doulames et al., 2023 show that injectable hydrogels with a stiffness matching the spinal cord and binding sites mimicking fibronectin binding domains enhanced iPSC neuronal integration with spinal cord tissue following a cervical injury[34]. However, compared to a biomaterial implant with no iPSC neurons or direct injection of iPSC neurons into the injury site in saline, no functional improvements in injured rats were observed compared to the combination of injectable hydrogels and iPSC neurons[34]. This lack of functional recovery may in part be due to the shear force of hydrogels passing through a needle for injection into the injury site, which in turn affects fragile iPSC neuron viability and is known to induce cell death[35,36]. Moreover, while our previous work showed that collagen-IV and fibronectin combinations influence whole-cell neural morphology in 2D and 3D environments[10,11,37], and that each molecule can modulate nanoscale actin architecture[32], here we demonstrate for the first time that this neurotrophic combination produces coordinated nanoscale and whole-cell changes in motor neurons, linking subcellular actin remodelling to enhanced neurite outgrowth. Therefore, the neurotrophic macroporous scaffolds presented in this study can be seeded and implanted directly to reduce iPSC neuronal death from shear forces[34–36].

To further enhance iPSC neuronal growth within the scaffold and promote integration of the delivered cells with injured tissue, we proposed to functionalize the soft, biomimetic scaffolds with genetic cargoes to promote neurite growth and overcome intrinsic barriers associated with neuronal repair. Through pathway enrichment analysis of publicly available datasets, we identify and show that miR-133a affects key downstream neuronal pathways, including those involved in actin regulation, neurotransmitter transport and synaptic vesicle cycling, all of which are dysregulated in injured neurons[38–40]. Mechanistic studies in the area of spinal cord and axonal regeneration have highlighted actin regulating pathways as key modulators of axonal extension following injury[2,25,41]. Studies investigating miR-133a activity in non-neural cells, such as bronchial smooth muscle cells, have also shown that miR-133a down-regulation also affects actin organisation, indicating that miR-133a and actin regulation are linked and these interactions are conserved across non-neuronal cells[27]. Therefore, miR133a inhibition was identified as a target that may promote neurite outgrowth through modulating actin regulation, a key process that is affected in the adult spinal cord following injury[2].

Non-viral miR-133a-i nanoparticles were successfully manufactured and, when characterised, showed optimal physicochemical properties for cell delivery with an N:P ratio of 8, and a suitable size, morphology, stability and charge for cellular internalisation[29]. When loaded into the biomimetic scaffolds, the miR-133a-i-activated scaffolds demonstrated sustained release, retaining >95% of the cargo for up to 28 days under static conditions *in vitro*. Initially, the scaffolds exhibited a burst release of genetic cargo within the first 24 h followed by a slower diffusion-mediated release over several weeks. While various factors such as nanoparticle size, binding domains and scaffold-cargo affinity can influence the release profile of gene cargo from scaffolds[42–44], our results align with previous studies using different vectors and cargos for different applications[20,29]. It is expected that the high nanoparticle retention may be due to our scaffold being comprised of hyaluronic acid which is negatively charged, which likely enhances the scaffold’s ability to absorb and retain positively charged nanoparticles via electrostatic interactions[45,46]. This retention is ideal for the transfection of the scaffold-seeded iPSC neurons, which were observed co-localising with the nanoparticles within the scaffold. Activated microglia/infiltrating macrophages in the injury site may also accelerate scaffold degradation and therefore enhance the rate at which nanoparticles are released into the external environment in an *in vivo* model of spinal cord injury[47,48]. Therefore, this passive release of gene-cargo is advantageous to respond to the phagocytic activity of activated glial cells while retaining a core portion of the scaffold to support neuronal delivery.

When seeded into the miR-133a-i-activated scaffolds, both motor neurons and iPSC neurons co-localised with nanoparticles, demonstrating efficient delivery of the loaded cargo directly to neurons. Direct gene and drug-based delivery to injured neural tissue can be challenging, as nanoparticles can slough away from the site of interest[16,49] and systemic delivery approaches face challenges such as crossing the blood-brain barrier[50]. By implanting a physiochemically matched scaffold that contains the gene cargo directly in contact with the neural tissue, both a supportive neurotrophic environment for cell growth is provided and more direct delivery of gene-cargo to the area of interest is facilitated. Additionally, the concentration of miR-133a-i used in this study did not exhibit cytotoxic effects on cells, as they fell within the optimal cytotoxic range (<3 μg) reported in previous literature[29,51]. Excellent viability and low levels of cytotoxicity as measured by metabolic activity and LDH release, respectively, were detected following delivery of miR-133a-i nanoparticles to both types of neurons, indicating that scaffold-based delivery can facilitate successful transfection to delicate neurons without compromising biocompatibility.

Bulk RNA-seq analysis provided mechanistic insight into how scaffold-mediated miR-133a inhibition enhances neurite outgrowth in iPSC-derived neurons. At the transcriptional level, miR-133a-i-activated scaffolds induced a coordinated upregulation of genes controlling actin dynamics, cell-matrix adhesion and vesicle trafficking, together with a broad increase in mitochondrial and oxidative phosphorylation pathways. For example, genes involved in Arp2/3-mediated actin branching and growth-cone motility[52], adhesion receptors[53], and regulators of vesicular transport[54] were enriched, suggestive of a cell-intrinsic program that favours cytoskeletal remodelling and membrane supply to elongating neurites[52,54]. In parallel, enhanced expression of respiratory chain components[55], mitochondrial enzymes[56] and translation factors[57] suggests that miR-133a inhibition shifts neurons toward a more energetically active state, ideal for supporting process extension and synaptic remodelling[55–57].

Conversely, miR-133a-i-activated scaffolds depleted a large cohort of mitotic[58], kinetochore[59], DNA-damage repair genes[60,61], and ciliary components[62,63], indicating a transition away from a proliferative or progenitor-like signature towards a more fully post-mitotic, projection-focused neuronal gene signature. Reduced expression of canonical mitotic regulators[64] and double-strand break repair factors, together with decreased levels of core centriole and intraflagellar transport genes[65], is consistent with prior work linking loss of these modules to terminal differentiation and neurite specialization[64]. Notably, multiple neuronal and synaptic genes were modulated, including adhesion molecules and pre-synaptic scaffold-related genes associated with synapse formation and maturation[66], implying that miR-133a-i-activated scaffolds may affect not only neurite elongation but also the assembly of nascent connectivity. Together, our data support miR-133a functions as an endogenous brake on a neurite-promoting, metabolically active neuronal phenotype where scaffold-mediated inhibition reprograms iPSC neurons towards a more permissive state for cytoskeletal re-modelling, outgrowth and integration with host tissue optimal for spinal cord repair.

The biomimetic miR-133a-i-activated scaffold supported the successful delivery of iPSC neurons to injured dorsal root ganglia explants, enhancing neurite extensions between the delivered cells and explants, while untreated scaffolds failed to do so. Neural repair strategies have struggled to successfully integrate delivered neurons into the injured tissue microenvironment due to inhibitory intrinsic[20,67] (i.e. inhibitory pathways in adult neurons preventing axonal extension) and extrinsic[68,69] (i.e. inhibitory ECM at the site of injury) factors. Superior cell engraftment is not defined only by cell survival but requires robust cell-cell interaction, appropriate positioning of transplanted neurons, and connectivity within host tissue[8,9,34,70]. While functional readouts remain the gold standard for measuring recovery of injured neural tissue *in vivo*, increasingly studies recognise that detailed morphological integration and scaffold-cell interface analysis predict longer-term functional outcomes[9,34]. In this study, we leveraged high-resolution imaging methodologies to investigate these phenomena at the scaffold-neural interface. Across untreated scaffolds, both DRG explants and iPSC neuronal spheroids maintained good viability and gross tissue contact. However, at higher resolution, cells mostly displayed relatively modest membrane extensions. These morphologies are characteristic of a less dynamic state, with relatively fewer cytoskeletal projections interacting with adjacent matrix or host tissue[71].

In contrast, within miR-133a-i-activated scaffolds, there was a notable increase in the complexity and extent of cell spreading, process extension, and substrate engagement, both at the scale of individual neurons and in overall tissue interface regions. Cells demonstrated cytoskeletal protrusions, more robust interaction with the scaffold, and broader intercellular contact. These morphological changes strongly suggest that scaffold-mediated inhibition of miR-133a liberates cytoskeletal reorganisation pathways, enhancing both the plasticity and adhesive interaction potential of the delivered neurons. Neuronal plasticity is considered crucial for axonal extension, synaptic targeting, and, ultimately, neural repair[71,72]. Confocal analysis paralleled these findings, showing improved spatial interdigitation of the border between host tissue and iPSC spheroid, implying more advanced stages of integration. These patterns mirror prior work in which successful axonal regeneration and neural engraftment were preceded by similar ultrastructural outcomes[9,11]. Overall, through a combinatorial, multifaceted approach of delivering neurotrophic gene cargo and providing a supportive scaffold, intrinsic and extrinsic barriers can be overcome to enhance neuronal delivery for spinal cord repair applications.

In addition to enhancing neuronal integration, miR-133a-i-activated scaffolds also appeared to engage neurovascular processes that may be advantageous for neural tissue repair. Transcriptomic profiling of iPSC-derived neurons in miR-133a-i-activated scaffolds revealed upregulation of several pro-angiogenic and adhesion-related genes[73], including secreted matricellular and growth-factor-like molecules[74–77], adhesion and junctional regulators[78], and immune modulators[79,80] linked to vascular remodelling. This profile suggests that neurons within miR-133a-i-activated scaffolds acquire a more angiogenic-competent genotype, with the potential to contribute to re-modelling the surrounding microenvironment in favour of vessel ingrowth and stabilisation. The induction of key neurovascular genes such as MFSD2A further hints at shared regulatory modules between neuronal miR-133a signalling and endothelial barrier or blood–brain barrier–associated pathways[81].

Functional assays in endothelial cells provided direct support for a pro-angiogenic influence of miR-133a inhibition. Treatment with miR-133a-i nanoparticles accelerated closure in scratch-wound assays promoted the formation of VE-cadherin-rich junctions and F-actin extensions spanning between cells, consistent with enhanced migratory capacity and network formation[82]. Importantly, these *in vitro* findings translated to an *in vivo* context using the chick chorioallantoic membrane model[83], where iPSC neuron-seeded miR-133a-i-activated scaffolds significantly increased average blood vessel branch length compared with untreated scaffolds. Collectively, these data indicate that the miR-133a-i-activated scaffolds exert dual neurotrophic and angiogenic actions by directly promoting neuronal outgrowth and integration with injured neural tissues, while simultaneously stimulating endothelial migration and blood vessel remodelling. Such coordinated neurovascular support is likely to be critical for long-term survival and functional integration of transplanted neurons in the hostile milieu of the injured spinal cord.

## Conclusion

In conclusion, this study demonstrates the successful development of a miR-133a-i-activated scaffold engineered to locally deliver nanoparticles to enhance iPSC-derived neuronal delivery and growth at the neural tissue interface while exploring the underlying mechanisms of action. Our transcriptomic data highlight coordinated changes in pathways associated with cytoskeletal remodelling, neuronal growth, and tissue repair, while complementary angiogenic analysis indicates that this miR-133a-i-activated scaffold also fosters a pro-vascular milieu that can further support spinal cord repair. Together, these findings position this dual-function, miR-activated scaffold as both a biomimetic neurotrophic substrate and an active regulator of neuro-regenerative and angiogenic responses, underscoring its potential as a next-generation platform for spinal cord injury and more broadly neural repair.

## Acknowledgements

This research was funded by a joint funding initiative of the Irish Rugby Football Union Charitable Trust (IRFU-CT) and the Advanced Materials and Bioengineering Research (AMBER) Centre through Taighde Éireann–Research Ireland (SFI/12/RC/2278). We also acknowledge funding from the Government of Ireland Higher Education Authority’s North-South Research Programme (FOB-HEANS-2022). R.M. received funding from a Research Ireland Government of Ireland PhD Studentship. M.D. received funding from a Research Ireland Government of Ireland Postdoctoral Fellowship. J.P. acknowledge funding from the European Union FET project PRIME- A Personalised Living Cell Synthetic Computing Circuit for Sensing and Treating Neurodegenerative Disorders’ (H2020 FET-GA 964712) and Taighde Éireann-Research Ireland (21/RC/10294_P2) FutureNeuro Research Centre for Translational Brain Science. Scanning electron microscopy was carried out using the Advanced Microscopy Laboratory (AML) facilities at Trinity College, Dublin. The authors acknowledge the support of Massimiliano Garre and the RCSI Super-Resolution Imaging Consortium funded by Taighde Éireann–Research Ireland (18/RI/5723). All raw and processed RNA seq data will be deposited in a public repository, and the accession number will be provided upon request. All other data that support the findings of this study are available from the corresponding author upon reasonable request. Tissue harvesting was approved by the RCSI Research Ethics Committee (REC202005013).

## Data availability statement

Data will be made available upon request.

## Materials & Methods

All reagents were purchased from Sigma–Aldrich (Ireland) unless stated otherwise. Wash steps refer to 3× Dulbecco’s phosphate-buffered saline (DPBS) washed for 5 min at room temperature (RT) unless stated otherwise. All cells were mycoplasma tested using a mycoplasma detection kit (Invivogen, USA). All tissue harvesting was approved by the RCSI Research Ethics Committee (REC202005013). All iPSC-derived cells were manufactured from a commercially available iPSC cell line obtained from Cedars Sinai Bio-manufacturing Centre, California, USA & the European Bank for iPSCs.

### ECM coverslip coating preparation

To assess the growth-supportive capacity of the neurotrophic ECM protein combination collagen-IV (CIV) and fibronectin (FN), glass coverslips were placed in 24-well culture plates and sterilised in 70% (v/v) ethanol (overnight, RT), followed by a DPBS wash. Coverslips were first coated with poly-L-lysine (PLL; control substrate, 10 μg/mL in DPBS, 1 h, RT), washed in DPBS, and then incubated with CIV and FN (each 10 μg/mL in DPBS, 1 h, RT). Finally, coverslips were washed again in DPBS before cell seeding.

### Scaffold manufacture

Hyaluronic acid (HyA) scaffolds of 3 mg/mL with and without CIV/FN were manufactured as previously described[10,11,33]. Briefly HyA sodium salt (1.6–1.8 MDa) and dH2O were added together to produce HyA solutions of 3 mg/mL and were mixed for 24 hrs. Next, 1-ethyl-3-(3-dimethylaminopropyl) carbodiimide (EDAC) (0.8 g/100 mL solution) and adipic acid dihydrazide (9.16 g/100 mL solution) were added in excess and allowed to mix for 24 hrs. 1 m hydrochloric acid was added to lower the HyA slurry pH to 4, initiating the crosslinking reaction (4°C, for 18 hrs). The pH was then raised to 7.4 using 1M sodium hydroxide and the crosslinked Hya was dialyzed for 6 days in sequential solutions of sodium chloride (NaCl) solution (29.5 g/L dH2O, 24 hrs, RT); fresh NaCl solution (29.5g/L dH2O, 24 hrs, RT); 20% ethanol (24 hrs, RT); dH2O replaced every 24 hrs (72 hrs total, RT). Collagen-IV and fibronectin were then added to the HyA solution (0.1 mg/mL each), syringe triturated (× 30 times) and then degassed to a pressure of 4000 mTorr (Leybold D16B, Biopharma, UK). To produce an individual scaffold, 250 µL of each slurry solution was pipetted into custom-designed PTFE molds coupled to metal baseplate using previously described protocols before being loaded into a freeze dryer (VirTis Advantage Pro, Biopharma, UK) set at −40°C for 25 hrs. Once completed the formed scaffolds were rehydrated in an ethanol series of 100% ethanol (1 hr, RT); 90% ethanol (1 hr, RT), and 70% ethanol (overnight, 4°C) followed by incubation in 14 mM EDAC and 5.5 mM N-hydroxysuccinate (NHS) in 70% ethanol (2 hrs, RT) to crosslink collagen-IV/fibronectin into the HyA before performing a final DPBS wash step to produce established soft (∼0.8kPa), macroporous scaffolds that have been characterised extensively in our previous studies[10,11,13,33]

### Pathway analysis of miR-133a related signalling in publicly available datasets

To identify miR-133a as a candidate target impacting neuronal cytoskeletal regulation and growth, we performed an in silico analysis of publicly available miRNA–mRNA interaction datasets. Predicted targets of human miR-133a-5p were retrieved from TargetScan, miRDB and StarBase using default score thresholds for conserved and high-confidence interactions. Target lists from each database were intersected to generate a consensus set of miR-133a targets from all three databases (112 genes). The intersecting gene set was submitted and analysed in the Database for Annotation, Visualization and Integrated Discovery (DAVID) for Gene Ontology (GO) biological process and Kyoto Encyclopaedia of Genes and Genomes (KEGG) pathway enrichment. Enrichment analyses were performed and significance was defined as an adjusted p-value (padj) of <0.05. Enriched GO terms and KEGG pathways related to neuronal function, actin cytoskeletal organisation and synaptic/vesicular processes were prioritised to guide subsequent experimental design and selection of miR-133a as a neurotrophic target for scaffold-based gene activation.

### MiR-133a inhibitor nanoparticle formulation and characterisation of size, zeta-potential, morphology and complexation efficiency

The miRIDIAN microRNA hsa-miR-133a-5p inhibitor (referred to as miR-133a-i; Dharmacon, UK) was combined with the RALA peptide (Biomatik, US) at an N:P ratio of 8, previously identified as optimal for cell delivery[29]. Complexes were allowed to form spontaneously for 30 min at RT following established protocols[29,31]. Nanocomplexes containing genetic cargo were then lyophilised in 2 mL glass vials using an Advantage Pro freeze dryer (SP Scientific, USA), with trehalose included as a cryoprotectant.

The mean hydrodynamic diameter and zeta potential of freshly prepared and reconstituted nanoparticles containing 0.5 μg microRNA in water was measured by dynamic light scattering (DLS) using a Zetasizer Nano ZS (Malvern Instruments, UK), with all measurements performed at RT. Nanoparticle morphology was examined by transmission electron microscopy (TEM): 10 μL of sample was applied to copper–carbon mesh grids (TAAB Laboratories, UK) for 3 min, air-dried overnight and stained with UranyLess (EMS, USA) for 3 min at RT. Grids were imaged on a JEM-1400 Plus TEM (JEOL, USA) operating at 80 kV.

Complexation efficiency was assessed using fluorescence spectroscopy and ion-exchange chromatography. For fluorescence quantification, 0.5 μg microRNA was formulated at an N:P ratio of 8 and unbound microRNA was measured with the Quant-iT™ Ribogreen Assay (Thermo Fisher Scientific, UK) on a plate reader (BioTek Instruments Inc., UK) according to the manufacturer’s instructions; complexation efficiency was calculated relative to naked microRNA controls. For ion-exchange chromatography (IEC), 0.5 g SP-Sephadex (Sigma-Aldrich, SPC25120, Germany) was swollen overnight in 10 mL 1 M NaCl at RT, then washed three times with 10 mL ultrapure water. Aliquots (20 μL) of naked miR-133a-i or miR-133a-i-RALA nanoparticles (≥20 [µg]/mL) were loaded onto the column and eluted with 3 mL ultrapure water, and collected fractions were analysed by UV–Vis spectroscopy.

### Nanoparticle scaffold loading efficiency and release kinetics

To assess scaffold loading efficiency and release profiles of miR-133a-i nanoparticles, HyA scaffolds were placed in 24-well plates and 10 μL of miR-133a-i nanoparticles (total RNA 1 μg per scaffold) was added to each scaffold side. Scaffolds were incubated for 30 min at 37 °C to allow nanoparticle attachment, then either (i) collected immediately for loading efficiency analysis or (ii) transferred to fresh wells for release studies. For release, scaffolds were incubated in 500 μL DPBS at 37 °C under static conditions. At defined time points up to 28 days, 200 μL of DPBS was removed and replaced with 200 μL fresh DPBS. Collected samples were incubated with 50 μL heparin (1 mg/mL) for 90 min at 37 °C to dissociate nanoparticles and release the miR-133a-i. Released miR-133a-i was quantified using the RiboGreen assay (Invitrogen, UK) on a plate reader (Tecan, UK), and data were expressed as cumulative release relative to the loaded dose.

### Neuronal cell culture

To study the effect of ECM proteins on neuronal growth and morphology, NSC-34 mouse motor neuron-like cells (ATCC, USA) were cultured on CIV/FN-coated substrates. Cells were expanded in growth medium (DMEM high glucose supplemented with 10% foetal bovine serum (FBS), 1% penicillin/streptomycin (P/S) and 1% L-glutamine) in T-175 flasks (37°C, 5% CO₂) and used between passages 7-13. Cultures were fed every 2-3 days until 80-90% confluent, then detached with 5 mL trypsin-EDTA (37 °C, 5 min), neutralised with an equal volume of medium and centrifuged (1,000 rpm, 5 min). Cells were resuspended and seeded onto CIV/FN-coated coverslips at 2×10³ cells/coverslip. After 24 h in growth medium to aid post-seeding survival, cultures were switched to differentiation medium (DMEM/F-12, 1% P/S, 1% L-glutamine, 1% FBS, 1% non-essential amino acids (NEAA) and 5 μL per 50 mL of 50 μM all-trans retinoic acid( for a further 7 days to promote neuronal phenotypes. For scaffold studies, NSC-34 cells were expanded as above and seeded onto HyA scaffolds at 1.25×10⁵ cells per scaffold side (30 min at 37 °C before flipping and seeding the opposite side). Neurons were cultured on scaffolds for up to 7 days in differentiation medium, after which conditioned supernatant was collected for cytotoxicity profiling, metabolic activity was assessed, and scaffolds were fixed in 4% paraformaldehyde (PFA; 30 min, RT) for microscopy and morphometric analysis.

Human iPSC lines (NAS2 line: donated by Dr Tilo Kunath’s laboratory[84]; CS29Ials-C9n1.ISOD7 line: obtained from Cedars Sinai Bio-manufacturing Centre, California, USA) were differentiated into cortical neurons using a modified version of the dual-SMAD inhibition protocol[85]. The CS29Ials-C9n1.ISOD7 line was used for all experiments with the exception of initial scaffold optimization experiments in Figure 1 where the NAS2 iPSC derived neurons were used. Human iPSCs were seeded on Geltrex^TM^-coated six-well plates (Thermo Fisher Scientific, 11612149) and cultured until they reached at least 80% confluency. 2 ml of Neural Induction Medium (NIM) was added to the cells. The NIM consisted of 2/3 (v/v) DMEM F12 with GlutaMAX™ (Thermo Fisher Scientific, USA; Cat. No. 31331093), 1/3 (v/v) Neurobasal plus Medium (Thermo Fisher Scientific, USA; A3582901), N2 supplement (50X) (Thermo Fisher Scientific, USA; 17502001), B27 plus supplement (100X) (Thermo Fisher Scientific, USA; A3582801) and 0.1 mM β-Mercaptoethanol (Gibco, Thermo Fisher Scientific, USA; 31350010) and GlutaMAX™ (Thermo Fisher Scientific, USA; 35050038). Media was supplemented daily with freshly prepared 100 nM LDN193189 (MedChemExpress, USA; HY-12071) and 10 µM SB431542 (MedChemExpress, USA; HY-10431). The cells underwent a half media change every day until day 10. On day 10, cells were passaged onto Geltrex^TM^-coated 12 well plates using 0.5 mM EDTA (0.5 M, pH 8.0, RNase-free; Invitrogen™, Thermo Fisher Scientific, USA; Cat. No. AM9260G). Half of the medium without small molecules was changed every other day until day 20.

Differentiated iPSC-derived neurons were maintained in neuronal maintenance medium (NMM; 50:50 DMEM/F-12 and Neurobasal Plus medium, 1% P/S, 1% GlutaMAX, 1% MEM NEAA, 0.2% N2 supplement, 1% B27 Plus and 50 μM 2-mercaptoethanol). Cells were passaged using Accutase (Gibco, UK), re-suspended in NMM and seeded onto HyA or HyA + CIV/FN scaffolds at ∼1.25×10⁵ cells per scaffold side (30 min at 37°C before flipping and seeding the other side). NMM was supplemented with 50 μM ROCK inhibitor Y-27632 dihydrochloride (Axon Medchem, Netherlands) before passaging or cell seeding to support survival, after which medium was replaced with fresh NMM and cultures were maintained for up to 7 days. At day 7, media were collected and immediately stored at −20 °C, and scaffolds were fixed in 4% PFA (30 min, RT), washed in DPBS and stored at 4 °C for downstream analyses.

### Neuronal immunostaining and image analysis

To analyse changes in neuronal morphology on ECM-coated substrates, neurons were stained, imaged and analysed as described previously[10,11,32]. Briefly, fixed NSC-34-seeded coverslips were immunostained for β-tubulin III (1:1,000; T3952, Sigma-Aldrich) followed by Alexa Fluor 555 goat anti-rabbit secondary antibody (1:500; Invitrogen, UK), and co-stained with Alexa Fluor 488 Phalloidin (1:1,000; Invitrogen, UK) and DAPI (1:500). Cells were imaged by confocal microscopy and reflection imaging on a Leica Stellaris 8-STED system (Leica, Germany). For nanoscale analysis of actin architecture, cells were stained with Alexa Fluor 594 Phalloidin (1:1,000; Invitrogen, UK) and imaged using a Leica STELLARIS 8 τ-STED 3D Falcon microscope with a 93×/1.30 glycerin immersion objective and laser settings as described previously[32]. For imaging of NSC-34 and iPSC neurons within scaffolds, samples were fixed and immunostained for β-tubulin III, Alexa Fluor 488 Phalloidin and DAPI using the same protocols as above[10,11]. Stained scaffolds were imaged on a Zeiss Examiner.Z1 confocal microscope (Zeiss, Germany) at 10× or 20× magnification, and representative z-stacks were converted to maximum-intensity projections. For iPSC neuronal spheroids, area, perimeter, Feret diameter (maximum width) and circularity, as well as NSC-34 neuronal area coverage, were quantified from thresholded images using ImageJ (Fiji). The number and distance of migrating cells and max neurite length per field of view were manually measured in ImageJ.

### Scanning electron microscopy of neuronal seeded scaffolds

For analysis of scaffold samples by scanning electron microscopy, fixed scaffolds were dehydrated as previously described[86], through an ethanol series (20%, 50%, 70% and 90% v/v; 10 min each at RT) and then stored in 100% ethanol overnight at RT. Samples subsequently underwent critical point drying in a critical point dryer (Quorum Technologies, UK) using liquid CO₂ with a bleed time of 1 h. Dried scaffolds were mounted on adhesive carbon stubs (Ted Pella, USA) and sputter-coated with a 5 nm layer of an 80:20 gold:palladium alloy using a Cressington 108 auto sputter coater. Images were acquired at varying magnifications on a Zeiss Ultra field-emission scanning electron microscopy (Zeiss, Germany) using the InLens detector (accelerating voltage of 5 kV), a 30 μm aperture and a 5 mm working distance.

### Metabolic, cytotoxic, cytokine release and glutamate uptake analysis

Metabolic activity was assessed using the Alamar Blue assay (Invitrogen, UK) according to the manufacturer’s instructions, by adding 300 μL reagent per well and incubating for 1 h at 37°C before fluorescence measurement. Lactate dehydrogenase (LDH) release, as an indicator of cytotoxicity, was quantified in conditioned media from scaffold cultures using the CyQUANT LDH cytotoxicity assay kit (Invitrogen, UK) following the supplier’s protocol. Glutamate uptake was evaluated in conditioned media using a fluorometric glutamate assay kit (Abcam, UK) according to the manufacturer’s instructions, and uptake was calculated relative to unconditioned control medium. For cytokine quantification, enzyme-linked immunosorbent assays (ELISAs) for mouse interleukin-1β (IL-1β; DY204, Bio-Techne, UK), tumour necrosis factor-α (TNFα; DY410, Bio-Techne, UK) and interleukin-10 (IL-10; DY217B, Bio-Techne, UK) were performed on conditioned media as per the manufacturer’s protocols. Absorbance was measured at 450 nm and 540 nm and cytokine concentrations were determined by interpolation from standard curves. All assays were performed using an Infinite® 200 PRO plate reader (Tecan, Switzerland).

### Bulk RNA-sequencing and transcriptomic analysis of iPSC neurons in miR-133a-i-activated scaffolds

For bulk RNA-sequencing of iPSC neurons cultured in untreated and miR-133a-i-activated scaffolds, scaffolds were harvested at day 3 for lysis and RNA extraction. Three scaffolds were pooled per sample in 500 μL QIAzol lysis reagent (Qiagen, UK), mechanically dissociated and immediately stored at −80 °C. Samples were subsequently thawed, and scaffold-free lysates were transferred to RNase-free tubes. Total RNA was isolated using the RNeasy kit (Qiagen, UK) according to the manufacturer’s instructions. Libraries were prepared and sequenced by Novogene (UK) using poly(A)-enriched mRNA libraries and a NovaSeq X Plus platform (paired-end 150 bp; N=4 untreated, N=5 miR-133a-i-activated). Reads were aligned to the human reference genome with STAR, and differential expression analysis was performed using DESeq2. Genes with an adjusted p-value (padj) <0.05 were considered significantly differentially expressed. Gene Ontology (GO) and Kyoto Encyclopaedia of Genes and Genomes (KEGG) pathway enrichment analyses were carried out by Novogene using standard pipelines.

### Analysis of miR-133a-i-activated scaffolds to enhance iPSC neuronal delivery to neural tissue explants

To assess whether iPSC neuronal-seeded miR-133a-i-activated scaffolds could promote neurite extension towards injured neural tissue, dorsal root ganglia (DRG) from 20-week-old wild-type C57BL/6 male mice were dissected and cultured on scaffolds as described previously[11,87]. All procedures complied with institutional animal ethics guidelines and the 3Rs principles. Briefly, DRGs were isolated using the hydraulic extrusion method[87], nerve roots were trimmed, and ganglia were immediately transferred to dissection medium (DMEM/F-12, 1% P/S, 1% L-glutamine, 25% heat-inactivated horse serum (Gibco, UK)).

Individual DRGs were gently placed onto untreated or miR-133a-i-activated HyA scaffolds that had been pre-seeded with iPSC-derived neurons for 7 days, and co-cultured for a further 7 days in iPSC NMM (as per *neuronal cell culture*). Explant cultures were fed every 48 h, and conditioned media collected at day 14 for glutamate uptake, ELISA and LDH release analyses. At day 14, DRG-scaffold cultures were fixed in 4% PFA (30 min, RT), washed in DPBS and stored at 4 °C until processing. Fixed samples were either (i) stained with Alexa Fluor 488 Phalloidin and DAPI (1:500; as per *Neuronal immunostaining and image analysis*) to visualise actin-rich processes and extensions between iPSC neuronal spheroids and DRG tissue, or (ii) processed for scanning electron microscopy via ethanol dehydration and critical point drying to examine ultrastructural features such as cell spreading and neurite extension at the DRG-scaffold interface (as described in *Scanning electron microscopy of neuronal seeded scaffolds*).

### Assessment of pro-angiogenic ability of miR-133a-i on endothelial cell migration and junction formation

To assess the capacity of miR-133a-i to enhance endothelial cell migration and junctional remodelling, a scratch-wound assay was performed as previously described[88]. Human umbilical vein endothelial cells (HUVECs; Lonza, Switzerland) were cultured in endothelial growth medium-2 (EGM-2) supplemented with SupplementMix (PromoCell, Germany). For migration assays, HUVECs were seeded in 12-well plates at 5×10⁴ cells/well and grown to confluence over 48 h. Monolayers were washed with DPBS and scratched using a sterile P1000 pipette tip to create a linear wound. Wells were rinsed to remove detached cells and then incubated in endothelial basal medium with or without freshly prepared miR-133a-i nanoparticles (1 µg miR-133a-i per well). Plates were transferred to a Zeiss Cell discoverer 7 microscope (Zeiss, Germany), and phase-contrast images were acquired every hour for 48 h. Scratch closure was quantified in ImageJ as the change in wound area relative to the initial time point (0 h). At the end of the assay, cells were fixed in 4% PFA (30 min, RT), permeabilised and immunostained for F-actin (Phalloidin) and VE-cadherin/CD-144 (diluted 1:200; 14-1449-82, Invitrogen, UK; as described in *Neuronal immunostaining and image analysis*) and imaged using an Examiner.Z1 confocal microscope (40× objective). Nuclei number and VE-cadherin-positive actin protrusion lengths per field of view were measured manually in ImageJ.

### Assessment of pro-angiogenic effect of miR-133a-i-activated scaffolds in an *in vivo* chicken chorioallantoic membrane model

To assess the angiogenic response to miR-133a-i-activated scaffolds *in vivo*, a chicken chorioallantoic membrane (CAM) assay was performed as previously described[88]. Fertilised chicken eggs (Shannon Vale Foods, Ireland) were cleaned and incubated at 37°C in a humidified incubator for 3 days. The eggshells were then carefully cracked to remove the upper shell, creating a window over the embryo, and embryos were incubated for a further 4 days. Untreated and miR-133a-i-activated HyA scaffolds pre-seeded with iPSC-derived neurons were gently placed onto the CAM, each positioned over a large blood vessel. Embryos were returned to the incubator and maintained for an additional 5 days. At the end of the incubation period, the vasculature surrounding each scaffold was imaged using a Nikon D5600 camera equipped with a Nikon AF-P DX Nikkor 18-55 mm f/3.5-5.6G lens at a fixed distance and magnification. Scaffolds were then carefully removed, and embryos were euthanised by decapitation and transection of central vasculature. Vascularisation was quantified on CAM regions surrounding each scaffold using an automated analysis script in ImageJ. Vessel length, branch number and junction number were measured and compared between untreated and miR-133a-i-activated scaffolds seeded with iPSC neurons.

### Statistical Analysis

Data and statistical analysis was carried out using Microsoft Excel and GraphPad Prism 8.0. All data were subjected to a Shapiro-Wilk’s normality test before applying appropriate parametric analysis. Where two treatments were compared, an unpaired, two-tailed *t*-test was used. Where more than one treatment was compared a one-way analysis of variance (ANOVA) with a Tukey post-hoc test was used. Where more than one treatment was compared across two factors, a two-way ANOVA with Bonferroni post-hoc test was used. All experiments were performed in triplicate. Results were expressed as mean ± standard error of the mean (SEM) except for neurovascular gene expression which was displayed using log2 fold change and adjusted p-values (padj). For all data *=*p*<0.05, **=*p*<0.01, ***=*p*<0.001 & ****=*p*<0.0001.

