## Supplementary Data for "Biomimetic miR-133a inhibitor activated scaffolds optimised for spinal cord repair promote neurite outgrowth and angiogenesis via neuronal cytoskeletal remodelling"

### Supplementary Figures

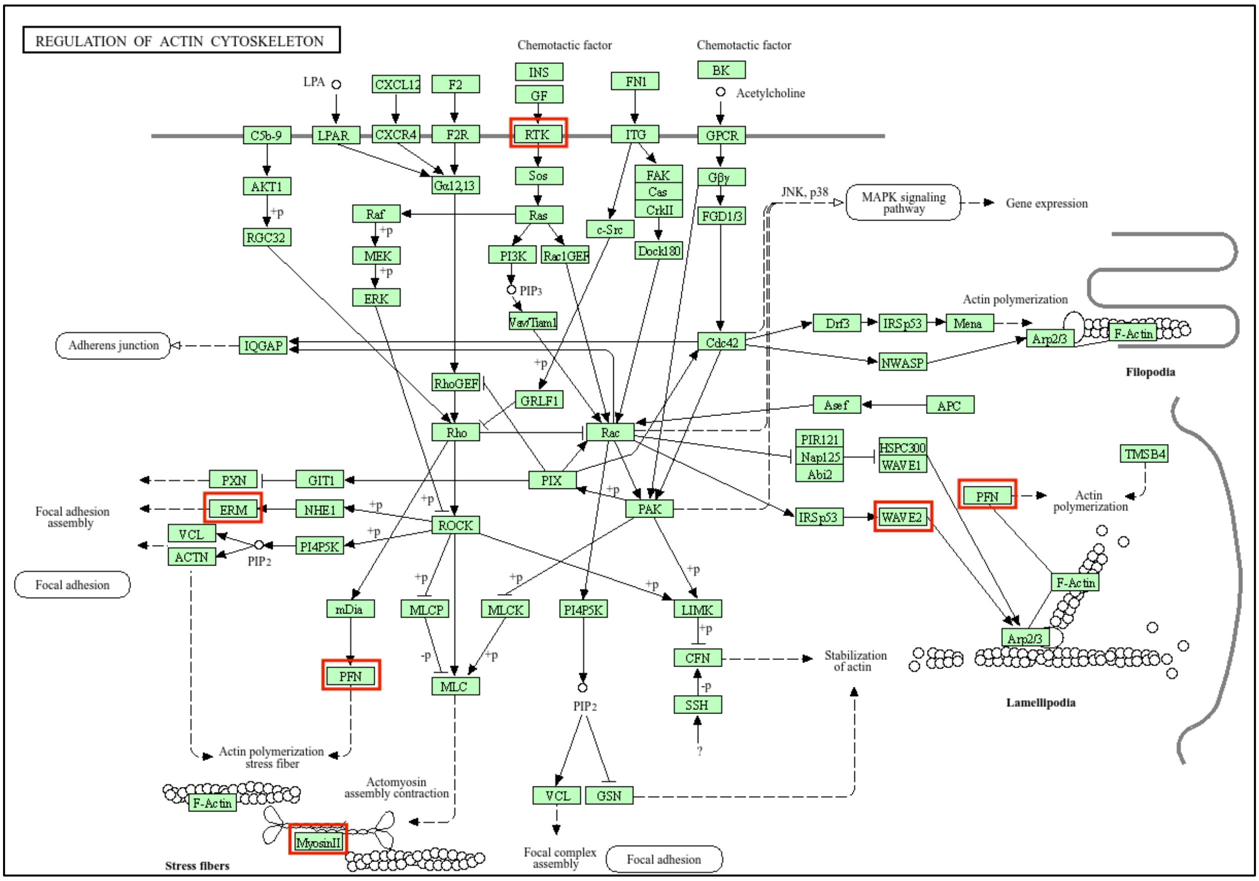


**Supplementary Figure 1. MiR-133a affects actin and cytoskeleton-regulating pathways**. Bioinformatics analysis of publicly available databases shows that miR-133a affects neuronal-related actin and cytoskeleton-regulating pathways.

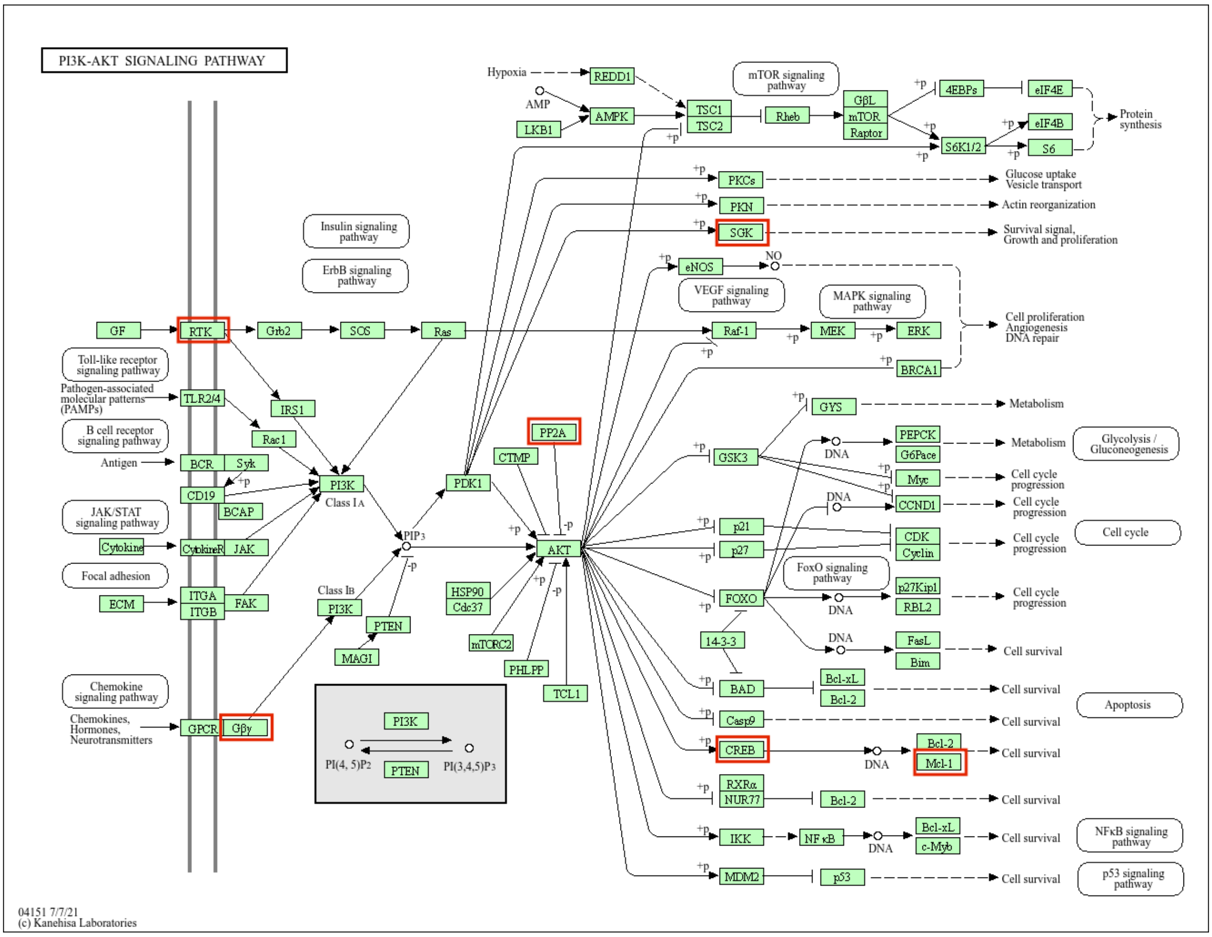


**Supplementary Figure 2. MiR-133a affects PI3K-AKT signalling pathways**. Bioinformatics analysis of publicly available databases shows that miR-133a affects neuronal-related PI3K-AKT signalling.

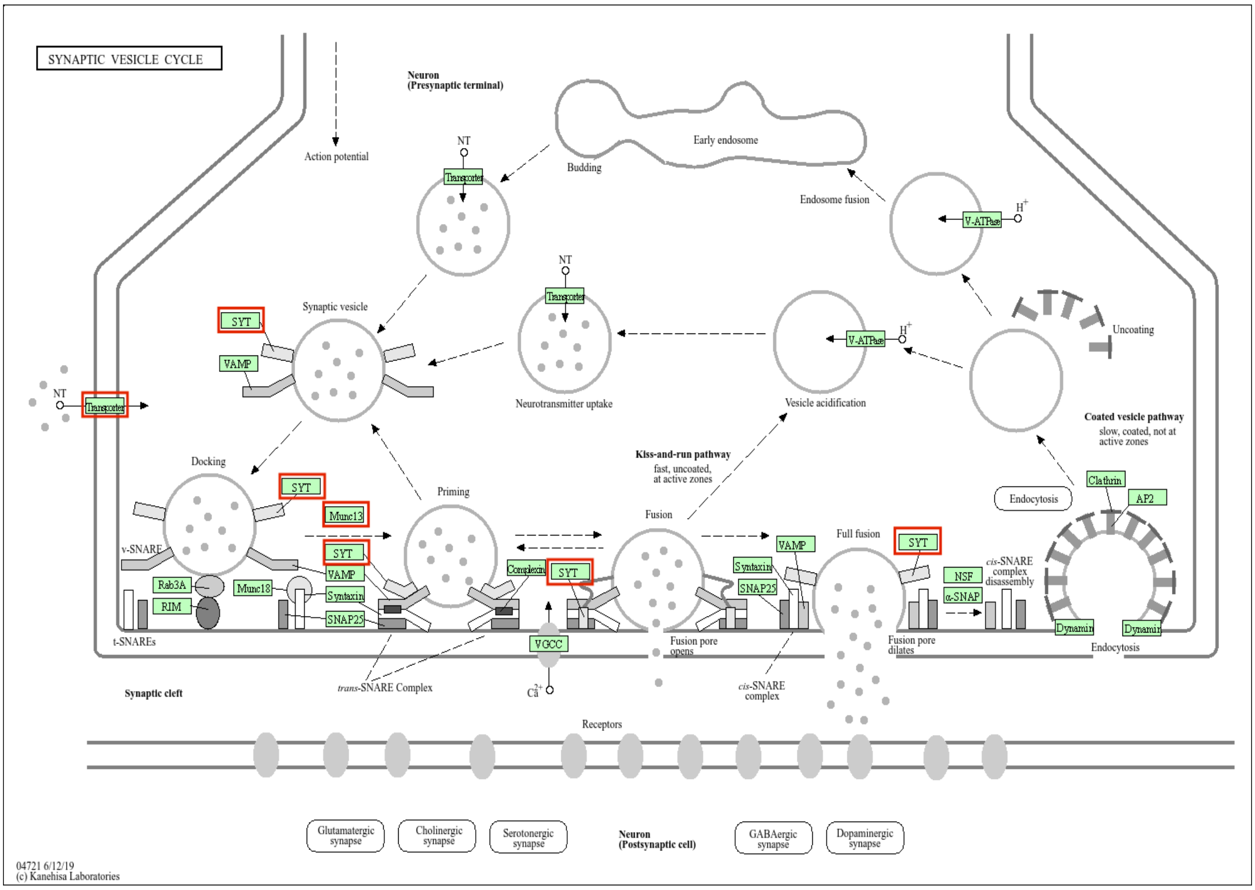


**Supplementary Figure 3. MiR-133a affects synaptic vesicle cycling signalling pathways**. Bioinformatics analysis of publicly available databases shows that miR-133a affects neuronal-related synaptic vesicle signalling.

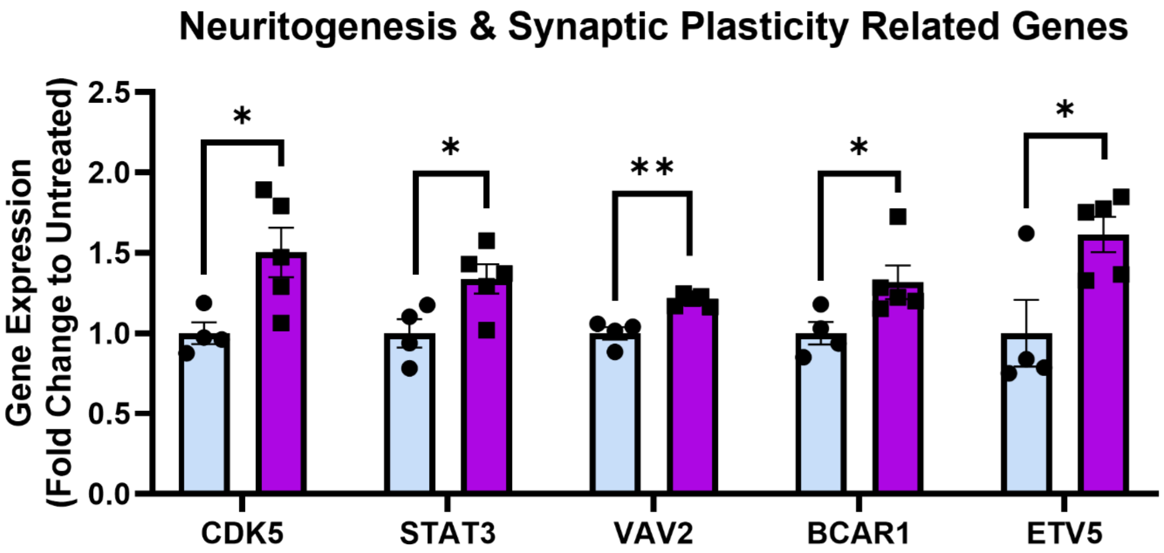


**Supplementary Figure 4. Transcriptomic profiling of iPSC neurons in miR-133a-i-activated scaffolds reveals modulation of neuritogenesis & synaptic plasticity-related genes.** Normalised iPSC neuronal gene expression changes for representative significantly upregulated genes involved in neuritogenesis and synaptic plasticity in untreated versus miR-133a-i-activated scaffolds. N=4-5. Analysis via an unpaired, two-tailed t-test for each respective gene. *p<0.05, **p<0.01.
